# Causal Discovery of Synchronous Neural Oscillations based on Jacobian-informed VAR-LiNGAM

**DOI:** 10.64898/2026.04.28.721377

**Authors:** Hiroshi Yokoyama, Ryosuke F. Takeuchi, Shohei Shimizu

## Abstract

The primary objective of system neuroscience is to understand the functional mapping and its causation in the dynamics of the brain network. Some experimental and methodological studies suggest that functional modularity and its hierarchical information processing in the brain network are crucial to understanding the functional role of task-specific or state-specific information flow in the brain. However, because most of the established techniques for detecting effective network structures in the neuroscience research field are strongly based on the “Granger causality” perspective, existing causal discovery methods specified for brain network analysis cannot identify the causal hierarchy in the modular network in the brain due to spurious correlation issues and indistinguishability of causal direction under the Gaussianity of observational noise in a linear system. To address the issues, we developed a causal discovery method for synchronous neural dynamics, called the Jacobian-informed linear non-Gaussian acyclic model, “j-VAR-LiNGAM”, by incorporating the information of the Jacobian matrix determined from a phase-coupled oscillator model estimated from observed neural data into the VAR-LiNGAM algorithms. The method was validated by showing that it could extract causal ordering in both synthetic data and empirical neural observed data. Moreover, by analyzing the observed neural oscillatory signals obtained from mice and humans, we confirmed that our method identified causally hierarchical structures in the brain, which aligned with the neurophysiological interpretations. These findings suggested that our proposed method can reveal the neural basis of hierarchical information processing in the brain network.

## 1 Introduction

Understanding the flow of information in the brain is crucial for revealing the functional role in brain network dynamics and neural mechanisms. The hierarchical information processing in the cerebral cortex of the brain is revealed in the anatomical evidence in the animal studies (Essen et al., 1992; Felleman & Essen, 1991). Moreover, recent neuroscience studies suggest that inter-regional communication and information processing in complex brain networks are inherently directional, organized by modular and hierarchical poly-synaptic signal propagation (Greaves et al., 2025; Park & Friston, 2013; Seguin et al., 2023). Although the complex brain network contains bidirectional inter-cortical feedback loops in anatomical brain connectivity (Essen et al., 1992; Felleman & Essen, 1991), these recent studies (Greaves et al., 2025; Park & Friston, 2013; Seguin et al., 2023) indicate the existence of task-dependent directionality in information processing within the networks. These findings support the notion that behind the complex network dynamics in the brain lies a structural information flow organized by a task-related or state-dependent core brain region (i.e., a causal source), which plays the role of an interconnected hub among functionally hierarchical network modules. In another context of studies on network neuroscience, researchers discuss the study policy and approaches to understand the causal and generative mechanisms of the functional role in complex network dynamics (Baker et al., 2022; Ross & Bassett, 2024; Siddiqi et al., 2022). However, most established causal analysis methods in network neuroscience are strongly dependent on the “Granger causality (GC)” perspective(Granger, 1980). For example, the directed transfer function (DTF), partial directed coherence (PDC)(Astolfi et al., 2008), and transfer entropy (TE)(Gourévitch & Eggermont, 2007; Schreiber, 2000), which are variations of the GC-type causal discovery (CD) method, contain the technical issues regarding the unidentifiability of causal direction and causal ordering in multivariate data. These issues come from the following two issues. The first issue is the definition of causality in GC perspective. In this perspective, the causality between two variables *Y* and *X* is based on the predictability. For example, if the predictability of the future state of *Y* is improved by adding covariates to the predictor with the past state of *X* as well as the past state of *Y*, the causation *X* → *Y* is established and vice versa. Thus, to establish the causality, the GC-based CD method requires temporal information and acceptance of a feedback loop (i.e., bidirectional causality) between variables. This issue would easily lead to the difficulty of identifiability in the causal ordering. The second issue is attributed to the Gaussian symmetry of the covariance matrix. Since both covariance Cov(*X, Y*) and Cov(*Y, X*) is similar due to Gaussian property, the linear Gaussian model (e.g., Vector auto-regressive model using linear GC test) cannot identically distinguish the causal direction *X* → *Y* or *Y* → *X* (Shimizu, 2014; Shimizu et al., 2006). Another approach of causal analysis in complex brain networks is the dynamic causal model (DCM) (Penny et al., 2004). This approach employs the state-space modeling (SSM) based model fitting into a dynamical system for observational data. Therefore, the latent dynamics of SSM formulated by the dynamical systems formulated as ordinary differential equations (ODEs) can be considered as the causally generative process of observations (i.e., the state equation explains the causal mechanism by which the observation is generated). Unlike the GC-based methods (DTF, PDC, and TE), the DCM is advanced enough to interpret the generative process and time-evolving dynamics in the neural data by applying SSM-based inference of a dynamical system (i.e., ODE). This allows us to interpret how the observed neural activity is generated and temporally evolved within the complex brain network. However, this approach is efficient only if the state of the dynamical model in the SSM is accurately inferred. Furthermore, because complex brain networks involve nonlinear interactions and bidirectional feedback loops (“cyclic” structures), it remains challenging to clearly identify causal order and hierarchy within these networks. Identifying causally hierarchical structures in the network requires determining the topological order of the estimated network graphs. In linear systems, this can be achieved by estimating a directed acyclic graph (DAG) with non-Gaussian noise using linear non-Gaussian acyclic model (LiNGAM) under the assumption of causal sufficiency (no unobserved confounders) (Shimizu et al., 2006). However, because brain network dynamics contain inherently nonlinear and time-evolving inter-cortical interactions, linear DAG-based approaches are difficult to directly apply to the neural data. Among nonlinear timeseries methods, the Peter and Clark Momentary Conditional Independence (PCMCI) approach (Runge et al., 2023) is well known; however, its reliance on PC-algorithm-based conditional independence tests limits the ability to uniquely identify topological order, as it cannot distinguish among graphs in a Markov equivalence class. Consequently, both GC and DAG-based methods face challenges in uniquely identifying causal hierarchies in complex brain networks from data.

In neuroscience, although many causal analysis methods have been developed to interpret directed and hierarchical communication in complex brain networks from data, most conventional approaches still struggle with unidentifiability when detecting causal mechanisms. To tackle this issue, we proposed a new causal discovery (CD) method enabling the estimation of the directed and hierarchical structures in nonlinear network dynamical systems such as complex brain networks. In our method, to consider both the dynamics of a time-evolving network and the identifiability of the causally directed information flow in the brain, we combined a data-driven modeling of dynamical network systems (i.e., ODE model) and a time-series causal discovery method. To establish such a method, we first introduced three assumptions: (1) synchronous oscillations in the neural data could be expressed as a network dynamical systems formulated as ODE like a phase-coupled oscillator model, (2) observations contain no unobserved confounders, and (3) directed “acyclic” causal structures (i.e., causal DAG structures) in the steady state of the ODE can be linearized as a structural vector auto-regressive (SVAR) model with Padé approximation under the non-Gaussianity assumption in observational noise. According to these assumptions, we developed a novel CD method specified network dynamical systems, incorporating the information of the Jacobian matrix in a phase-coupled oscillator model estimated from observed neural data (Yokoyama & Kitajo, 2022) into the vector-autoregressive LiNGAM (VAR-LiNGAM) algorithms for estimating SVAR model (Hyvärinen et al., 2010; Shimizu, 2014) (hereafter called j-VAR-LiNGAM: Jacobian informed VAR-LiNGAM). This enables us to identify the causal order in complex brain networks by extracting both instantaneous and lagged directional causal pathways reflected in the time-evolving dynamics of observed synchronous neural data.

To test the validity of our proposed method, j-VAR-LiNGAM, we applied this method to both numerical simulations and empirical neural oscillatory signals. In the numerical simulation, the method was applied to synthetic data generated by phase-coupled oscillators with known network structures. By doing so, we confirmed that our method can correctly detect causal structures and causal order in the network dynamical systems under the conditions that the systems exhibit weakly coupled limit cycle states. We then employ our method on two different datasets: laminar local field potential (LFP) that we obtained from in-vivo mice brain and an open dataset of human scalp electroencephalographic (EEG) signals recorded during epileptic seizures (Guttag, 2010; Shoeb, 2009). In this way, we tested whether our proposed method, j-VAR-LiNGAM, can be applied to detect the directed neural information flow and its underlying mechanisms in both the mesoscale local neural circuit (i.e., laminar LFP) and macroscale whole brain network (i.e., EEG).

## 2 Methods

In this section, we will describe the mathematical details of our proposed method and the validation procedures for the method. We first provide an overview of our proposed method and related mathematics. After doing so, we explain validation procedures using numerical synthetic data and empirical neurophysiological data.

### 2.1 Outline of our proposed method

In our study, we estimate the causal effect and causal ordering in a complex brain network by using a linearized time-series causal model in a data-driven manner. We note that we focus on the synchronous dynamics of neural oscillations in specific frequency bands (e.g., theta, alpha, and others) to estimate the causal structures in complex brain networks. However, as we mentioned in the Introduction section, the observed data obtained from the brain network systems contains the nonlinear time-evolving dynamics that would be explained as network dynamical systems formulated as an ODE, like a phase-coupled oscillator. Therefore, even though various cutting edging methods for CD are proposed in the context of statistical causal inference studies, conventional CD methods are difficult to apply directly to such observed data. To address this issue, we proposed the novel CD method that consists of the following three steps: (1) data-driven modeling of the dynamical network systems (i.e., estimating the model parameters in ODE formula in the systems), (2) extracting directed information pathway explaining as causal DAG from ODE systems using our proposed method (as describe later), and (3) evaluation of causal effect and its causal ordering to determining causal hierarchical structures in the systems (Fig. 1A). In the following section, we present detailed explanations for each three step procedure and describe how these procedures are linked to function cohesively as an integrated system.

**Figure 1:**
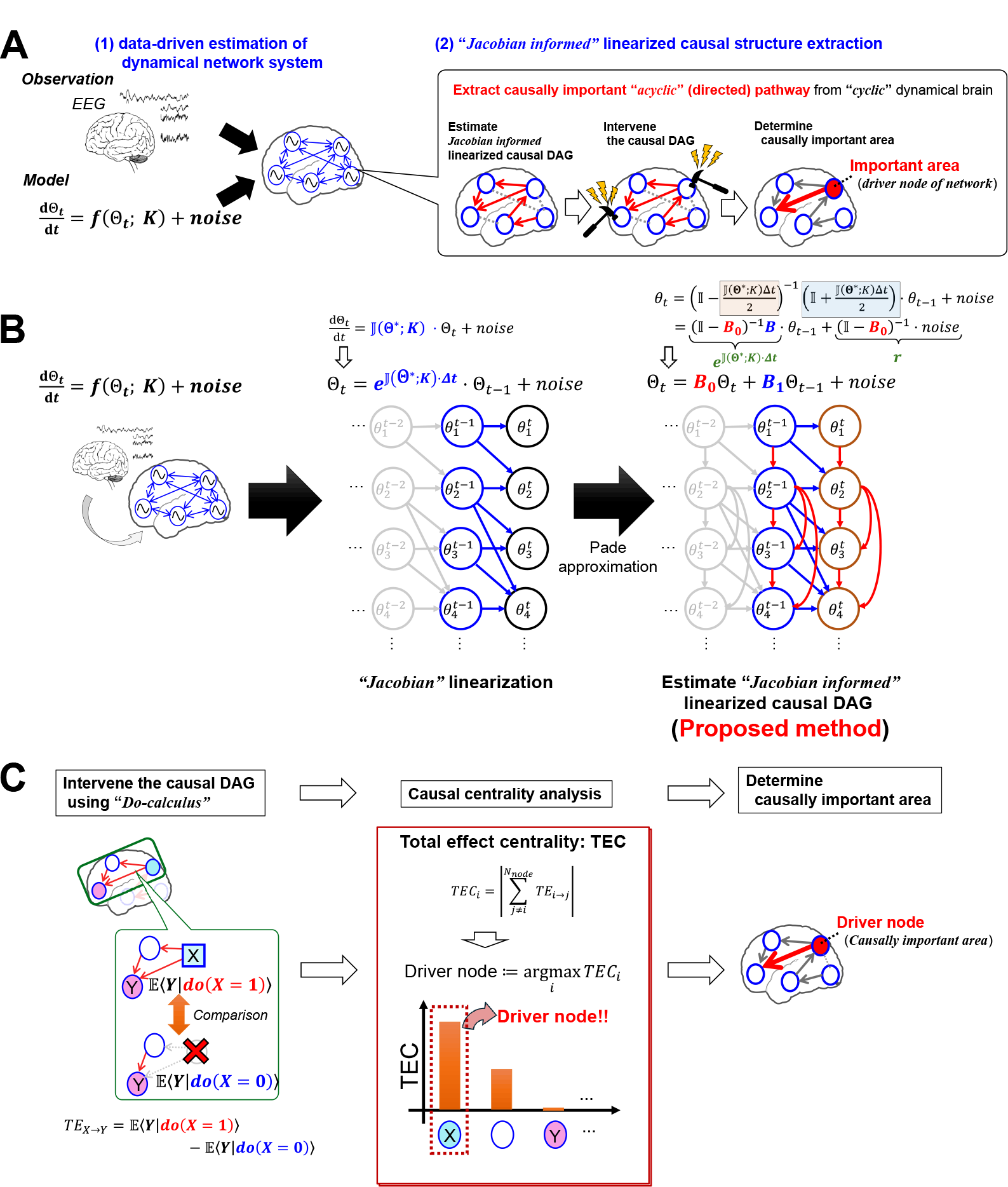
Outline of the proposed method, j-VAR-LiNGAM, for dynamical causal discovery. (A) Overview of the proposed method. Our method combines data-driven dynamical network modeling with time-series causal discovery. The central idea is to transform the estimated network dynamical model (which is a cyclic graph) into a time-series causal directed acyclic graph (DAG). This enables us to identify the causally significant regions (i.e., driver nodes). (B) Extracting causal DAGs from dynamical brain networks using j-VAR-LiNGAM. First, we estimate a phase-coupled oscillator model and its parameters in a data-driven manner. Next, we determine the Jacobian matrix and linearize the model using it. Finally, we convert the linearized model into a time-series causal model using j-VAR-LiNGAM, which incorporates the Jacobian matrix into the VAR-LiNGAM algorithm. For more details, please refer to the main text. (C) Estimating the driver node by using total effect centrality (TEC). We first calculate the total effect (TE) from selected node *i* to all nodes *j* that have node *i* as their ancestor, and sum up these TE values. The absolute value of this summation is defined as total effect centrality (TEC). After calculating the TEC for each network node, we identify the node with the highest TEC value as the driver node in that network.

**Figure 2:**
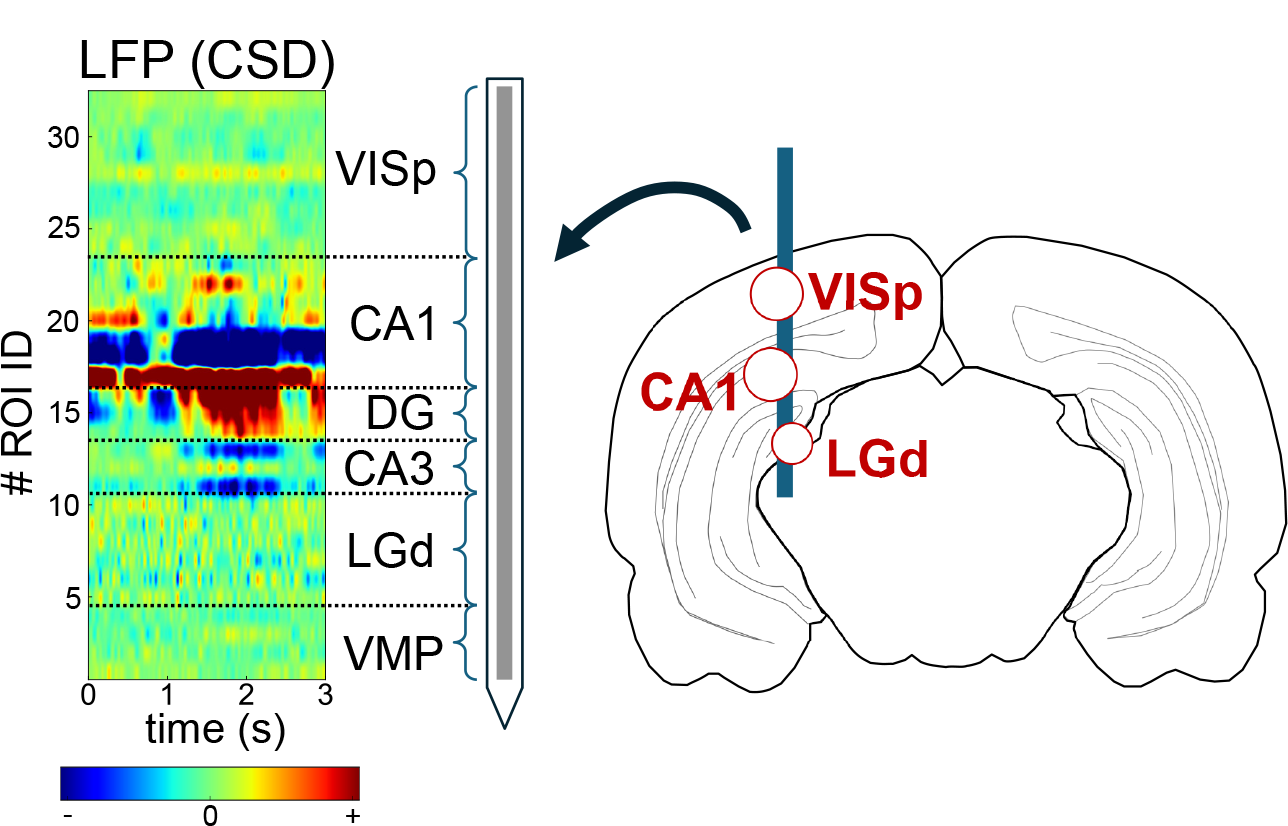
Schematic illustration of the laminar LFP recordings and its target brain regions. The LFP probe was inserted 4.0 mm below the cortical surface to measure the LFPs in primary visual cortex (VISp), hippocampus (CA1) and dorsal lateral geniculate nucleus (LGd). The left panel of this figure shows the schematic view of the laminar LFP observations (current source density) for each ROI (the number of ROI: 32) obtained from the mouse brain using a recording probe. The abbreviation of ROI labels indicates as follows: VISp: primary visual cortex, CA1, CA3: the first and third regions in the hippocampal circuit, DG: dentate gyrus, LGd: dorsolateral geniculate nucleus, and VMP: ventral midbrain*/*pons. Illustration of the mouse brain silhouette was sourced from Sci-Draw (https://scidraw.io/drawing/646).

#### 2.1.1 Step 1: Data-driven modeling of network dynamical systems

The first step is employed to determine the dynamical model in a data-driven manner. In this study, we assumed that the network dynamics reflected in the observed neural oscillations can be considered as a phase-coupled oscillator, and such an oscillator model is determined from observed data by using our previously proposed method (Yokoyama & Kitajo, 2022). Please refer to the original paper (Kuramoto, 1984; Netoff et al., 2012; Onojima et al., 2018; Ota & Aoyagi, 2014; Ota et al., 2020; Suzuki et al., 2018; Yokoyama & Kitajo, 2022) for details of the estimation procedures. However, we will here provide a brief summary of the outline of these procedures in Yokoyama and Kitajo (2022).

As mentioned above, for our method, we consider phase-coupled oscillators as the underlying time-evolving dynamics in complex brain networks. In that case, the temporal dynamics of *N* -th observed neural oscillations can be assumed to describe as the phase-coupled oscillator model using the following equation (see the previous works (Yokoyama & Kitajo, 2022) for a detailed description of the theoretical background):

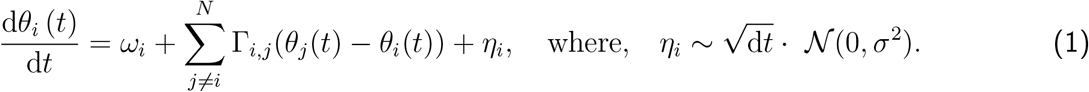

Note that *θ*_*i*_(*t*), *ω*_*i*_, and *η*_*i*_ indicate the instantaneous phase, the natural frequency, and the observation noise of the *i*-th oscillator, respectively. In our study, as in some computational and methodological neuroscience studies (Netoff et al., 2012; Onojima et al., 2018; Ota & Aoyagi, 2014; Ota et al., 2020; Suzuki et al., 2018; Yokoyama & Kitajo, 2022), given that the function Γ_*i,j*_ (*θ*_*j*_(*t*) − *θ*_*i*_(*t*)) is a 2-*π* periodic function when there exists a limit-cycle attractor of the dynamical system, we approximated the function 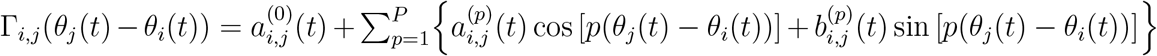 to the *P* -th Fourier series. As a result, Eq. (1) can be rewritten as follows.

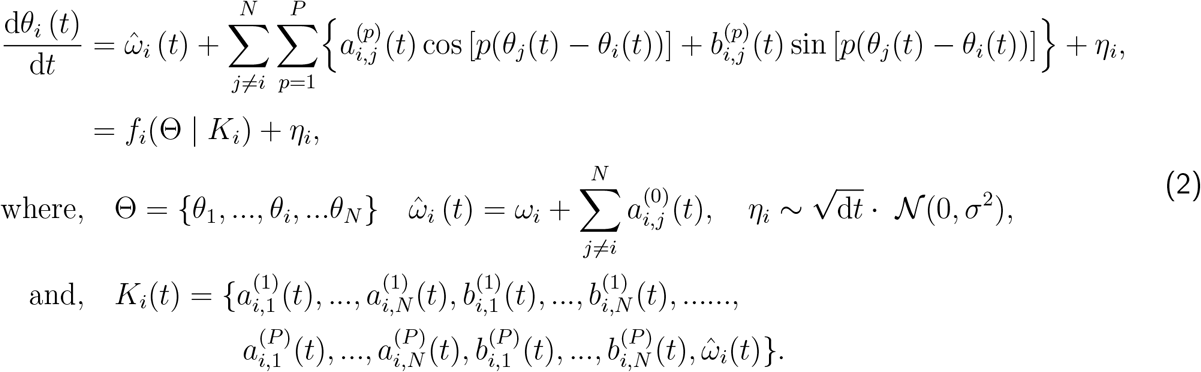

Given the prior distribution of the model parameters, *K*_*i*_ in Eq. (2) can be calculated using the Bayesian rule (Bishop, 2007; Sarris, 1973), as follows:

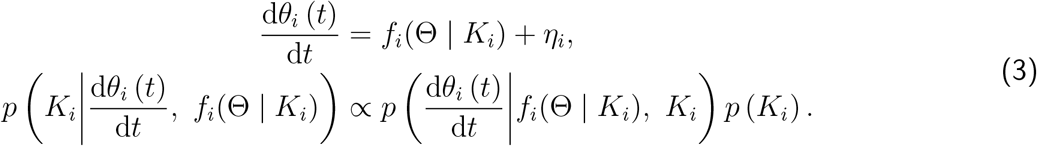

Here, in our proposed method, it is assumed that the observed noise *η*_*i*_ and prior to the parameter *K*_*i*_ have multivariate normal distributions, *η*_*i*_ ∼ 𝒩 (0, d*t* · *β*^−1^*I*) and 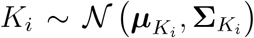, respectively. The likelihood of prediction: 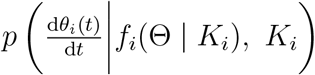 also follows the multivariate normal distribution 𝒩 (*f*_*i*_(Θ | *K*_*i*_), d*t* · *β*^−1^*I*). The parameter *β* is a precision factor of the observation noise that was fixed as an arbitrary value (e.g., *β*^−1^ = 0.1). The value of 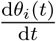 is approximated by the numerical differential: 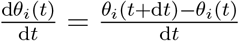. See Yokoyama and Kitajo (2022) for more details.

By solving the above Bayesian theorem in Eq.(3) to minimize the prediction error of the phase velocity 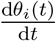, we can obtain the model statistics: 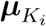 and 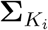 in the parameters *K*_*i*_ in a data-driven manner. The above procedures are illustrated on the left side in Fig. 1A, B.

For interconnecting the estimated phase-coupled oscillator (given as the ODE model) to the next step (described in the next section), we must determine the Jacobian matrix of the estimated ODE of the phase-coupled oscillator with fixed point solution for each phase 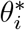. However, in non-equilibrium dynamical systems such as the brain (Nartallo-Kaluarachchi et al., 2026), it is difficult to estimate the fixed point solution from the ODE model obtained from a data-driven manner. Therefore, instead of the fixed point solution for each phase *θ*_*i*_ in the estimated phase-coupled oscillator model, we use the steady state solution of the phase difference Ψ_*i,j*_ = *θ*_*j*_ − *θ*_*i*_ in the model, which can be obtained from stationary distribution of the 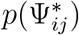 by solving the following Fokker-Planck equation of the model (Eq.(4)) (Ota et al., 2020).

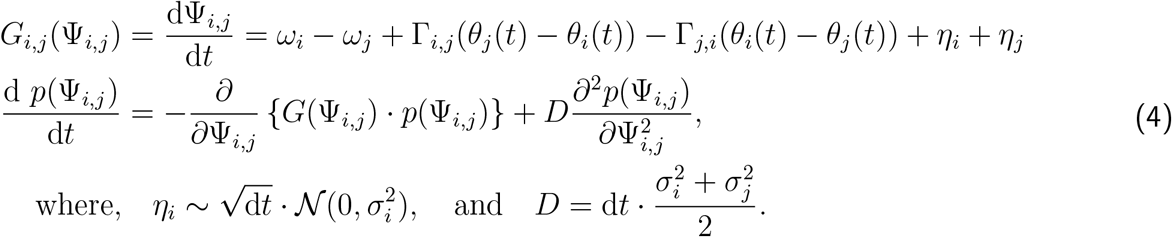

Here, *p*(Ψ_*i,j*_) stands the stationary distribution of the phase difference Ψ_*i,j*_ = *θ*_*i*_ − *θ*_*j*_ between the *i*- and *j*-th oscillators. By using Eq.(4), we can calculate the stationary distribution *p*(Ψ_*i,j*_) of the phase difference Ψ_*i,j*_ by numerically solving this equation until the distribution *p*(Ψ_*i,j*_) converges (Ota et al., 2020). In this study, the steady-state solution 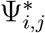 is defined as the value associated with peak of the distribution *p*(Ψ_*i,j*_), and this solution 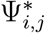 are calculated for all pair of oscillator *i, j*. As a result, the vector of all solutions 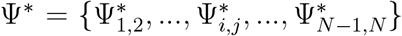 are adopted to estimating the Jacobian matrix 𝕁?(Ψ^∗^ | *K*) of phase coupled oscillators, which is formulated below.

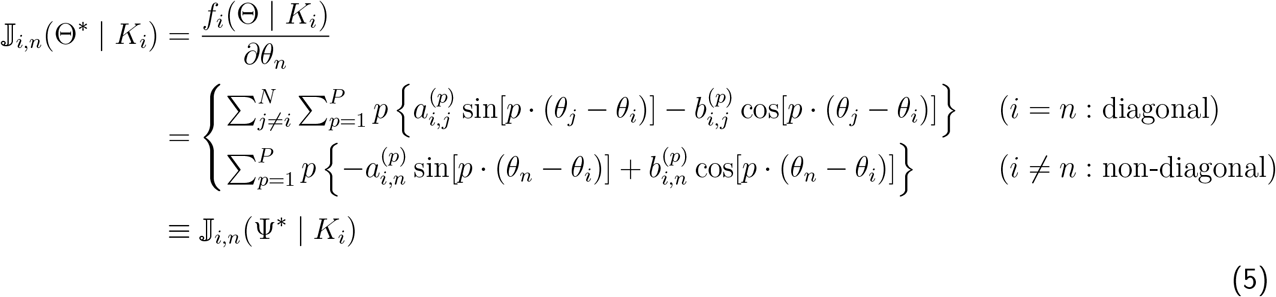

As can be seen in Eq.(5), each element of Jacobian matrix, 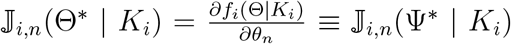, can be considered as the function of phase difference Ψ_*i,j*_. Therefore, if we can obtain the parameter and the steady-state solution of the Ψ by solving Eqs. (3) and (4), the Jaconian matrix of the phase-coupled oscillator can be calculated as the function of the Ψ^∗^ (i.e., 𝕁_*i,n*_(Θ^∗^ | *K*_*i*_) ≡ 𝕁_*i,n*_(Ψ^∗^ | *K*_*i*_)), without estimating the fixed point solution for each phase, Θ^∗^.

#### 2.1.2 Step 2: Transforming the dynamical systems to a causal model

The second step is adopted for transforming from an estimated network dynamical system (ODE, containing the “cyclic” structures) to Jacobian informed linearized causal model (containing “acyclic” structures). In this section, for ease of understanding the details of our proposed method, we first describe the definition and estimation algorithm of VAR-LiNGAM. Next, to bridge the phase-coupled oscillator and VAR-LiNGAM, we explained the relationship between VAR-LiNGAM and linearization of ODE using Padé approximation(Bergstrom, 1984; Oud et al., 2018; Oud & Delsing, 2010; Voelkle & Oud, 2013). Finally, we will provide the mathematical details and estimation algorithm of our proposed method, “j-VAR-LiNGAM: Jacobian-informed VAR-LiNGAM”. This method allows us to extract causal structures formed with DAG from priorly estimated ODE model (i.e., phase-coupled oscillator model). This section provides the key idea of our proposed method, illustrated in Fig.1B.

##### VAR-LiNGAM

As mentioned in the Introduction section, VAR-LiNGAM is the method to estimate the SVAR models with non-Gaussianity of the observed noise. The SVAR model is defined as follows.

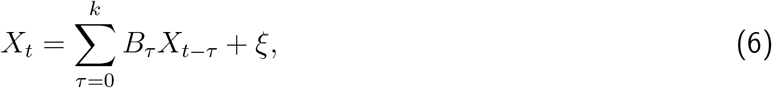

where *k* is the number of time-lags, indicating the model order of the SVAR model. *B*_*tau*_ is *N* ×*N* matrix of the linear coefficient at the time-lag *τ*. *ξ* denotes the observed noise. The most different point of the ordinary VAR model relative to the SVAR model is that the SVAR model considers the instantaneous effect *B*_0_ (*τ* = 0) for modeling the dynamics of the observed data. Moreover, by considering the following assumptions, we can obtain the instantaneous effect *B*_0_, represented with an “acyclic” graph, as is a typical causal DAG (Hyvärinen et al., 2010; Shimizu, 2014; Shimizu et al., 2006).

- The noise *ξ* are mutually independent
- The noise *ξ* follows non-Gaussian distribution
- The diagonal of instantaneous effect *B*_0_ is defined to be zero

We will then provide the estimation algorithm of SVAR model using VAR-LiNGAM. For ease of explanation, we describe the procedures of the estimation algorithm in VAR-LiNGAM with the first-order lag (i.e, the maximun lag *k* = 1). In that case, the Eq.(6) can be rewritten with *k* = 1 as follows.

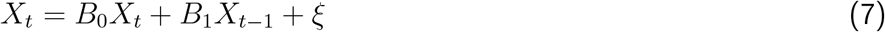

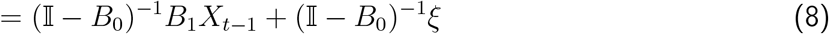

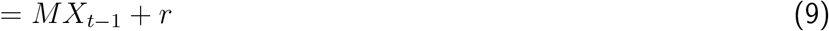

where *M* and *r* indicate the linear coefficient and the residuals in the ordinary VAR model. As can be seen in Eqs. (7–9), we can derive from the SVAR model to the ordinary VAR model form. In addition, considering the Eq. (8) = Eq. (9), the residual term *r* in Eq. (9) can be rewritten as follows.

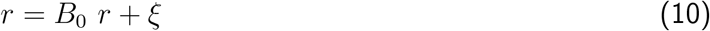

Moreover, if the Eq. (7) = Eq. (9), the residual *r* satisfy the following equivalence.

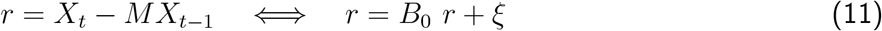

Thus, the *B*_0_ in Eq.(11) is identifiable as an “acyclic” graph under the non-Gaussianity assumptions in the noise *ξ*, by applying LiNGAM algorithms (Shimizu et al., 2006). If we obtained *B*_0_, because of Eq.(8) = Eq.(9), *B* is also predictable by using *B*_0_ and *M* .

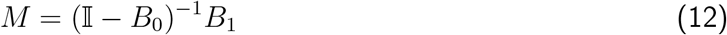

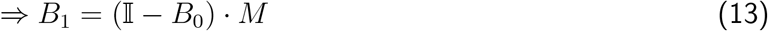

That’s the estimation procedures of SVAR model as causal model using VAR-LiNGAM. By using above method, we can uniquely identify the time-series causal model under the non-Gaussianity assumption for the observational noise *ξ* (Hyvärinen et al., 2010; Shimizu et al., 2006). However, because SVAR model assumed the linearity and weak-stationarity for the model, this model cannot directly applied to nonlinear time-series data generated by dynamical systems as the brain.

To address this issue, in this study, we proposed the method to bridge the dynamical systems (formulated by ODE) to SVAR model. For the following paragraph, we will describe how to link ODE with SVAR model.

##### Connecting from the ODE to the SVAR model

As described above, the VAR-LiNGAM method can be applicable for linear time-series modeling. Therefore, it is difficult to directly connect the ODE and SVAR models. However, by combining the Jacobian linearization and Padé approximation(Bergstrom, 1984; Oud et al., 2018; Oud & Delsing, 2010; Voelkle & Oud, 2013), we found that the solution of the linearized ODE can be approximated in a similar form to the first-order SVAR model (SVAR(1)). The details are described as follows.

Applying the Jacobian linearization, the solution of the nonlinear ODE can be obtained as follows.

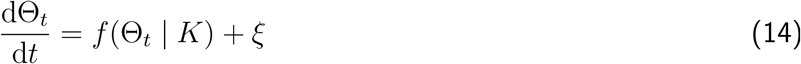

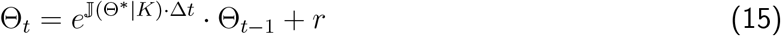

where, 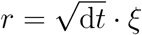 indicates the residual term.

By introducing the first order Padé approximation (Bergstrom, 1984; Oud et al., 2018; Oud & Delsing, 2010; Voelkle & Oud, 2013) instead of the Taylor approximation, the matrix exponent *e*^𝕁?(Θ ∗|*K*)*·*Δ*t*^ in the Eq.(15) can be expressed as the following approximated form.

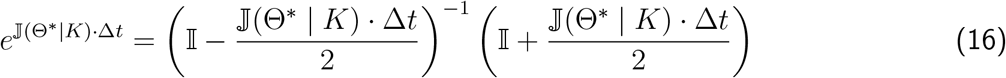

By inserting Eq.(16) into Eq. (15), the solution of the linearized ODE is given as the following equation.

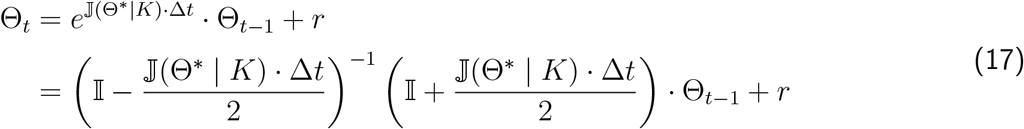

In that case, suppose 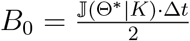 and 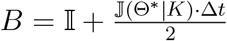, we can rewrite Eq. (17) as follows.

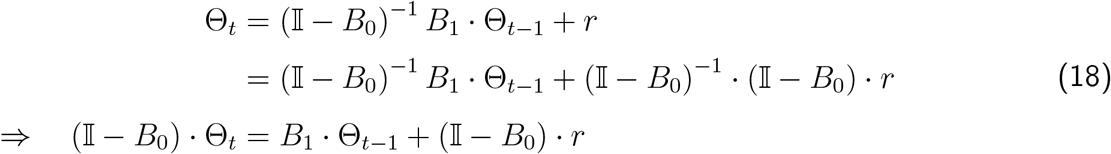

Therefore, Eq.(18) is finally rewritten as follows.

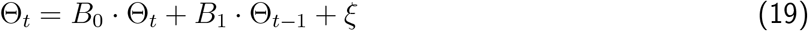

where, *ξ* = (𝕀 − *B*_0_) · *r*.

As can be seen in Eq.(19), this resulting equation is similar to the SVAR model. However, to consider Eq.(19) as a causal model, the instantaneous effect *B*_0_ should satisfy the DAG structures with the non-Gaussianity assumption regarding the noise term *ξ* in Eq.(19).

To do so, we proposed the following estimation algorithm inspired from the concept combining VAR-LiNGAM algorithm and linearized ODE solution with Padé approximation. In our proposed method, we first rewrite Eq. (19) in the same manner as Eqs. (7–9) as follows.

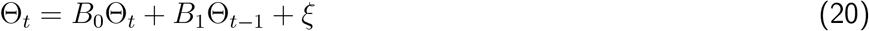

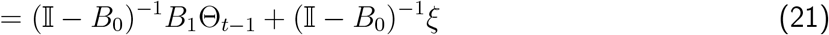

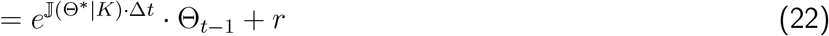

where, *r* = (𝕀 − *B*_0_)^−1^ · *ξ*.

According to the same manner in the VAR-LiNGAM algorithm, the residual *r* in Eq. (22) should satisfy the following relationship.

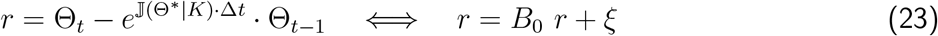

Therefore, if we can obtain the Jacobian matrix 𝕁(Θ^∗^ | *K*) of the ODE with the solution Θ^∗^ and a certain known parameter *K*, we can solve the instantaneous effect *B*_0_ in Eq. (23) by using VAR-LiNGAM.

Similarly, for the lagged effect *B*_1_ in Eq.(19), if the *B*_0_ is provided, the *B*_1_ can also be calculated as follows in the same manner in the Eqs. (12–13).

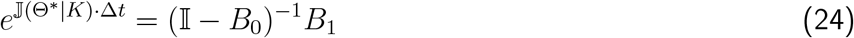

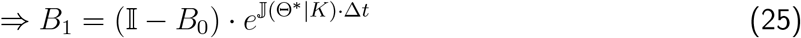

In summary so far, by utilizing Eqs. (15–25), we proposed the method to extract directed “acyclic” structures in the nonlinear ODE, which can be considered as a typical SVAR-based time-series causal model. In our proposed method, we first estimate the parameters of the phase-coupled oscillator using Step 1 outlined in section 2.1.1. Then, we linearize the ODE based on the estimated phase-coupled oscillator model using Padé approximation (Eq. (17)). After this, we apply Eqs. (23–25) to derive the causal DAG structures from network dynamical systems within the previously estimated phase-coupled oscillator model, employing a data-driven approach. We refer to the above proposed method as “j-VAR-LiNGAM (Jacobian-informed VAR-LiNGAM)”.

#### 2.1.3 Step 3: Determining causal ordering and hierarchically important area

As explained so far, we provided the key idea of our proposed method, j-VAR-LiNGAM, which allows us to extract causal structures from the estimated phase-coupled oscillator model by combining data-driven modeling of dynamical systems (Yokoyama & Kitajo, 2022) and VAR-LiNGAM-based time-series CD method (Hyvärinen et al., 2010; Shimizu, 2014). In this section, we provide the method to determine the causally important regions in the estimated causal model obtained from our proposed j-VAR-LiNGAM method based on the typical causal effect estimation approach with intervention (Bollen, 1987; Pearl, 2009a, 2009b).

By utilizing our method described in sections 2.1.1 and 2.1.2, we can derive the linearized time-series causal model, which is articulated as the Jacobian-informed SVAR (j-SVAR) model. Therefore, we can employ the total effect estimation using do-calculus (Bollen, 1987; Pearl, 2009a). At first, we shortly explain the outline of total effect estimation in the linear causal model, which is represented as a linear structural equation model (SEM). We here consider the linear SEM for *N* variables, parameterized by *N* × *N* matrix *B* ∈ ℝ^*N ×N*^ denoting the causal effect, as follows.

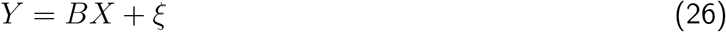

where *ξ* is mutually independent noise term. In this case, the total effect (TE) of the variables *X* on *Y* can be defined below (Bollen, 1987; Pearl, 2009a, 2009b).

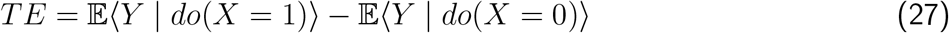

This quantifies the extent of changes in the outcome *Y*, reflecting the interventional effect between values by fixing *X* at *do*(*X* = 1) and *do*(*X* = 0).

This framework can be extended to a linear time-series causal model, such as the j-SVAR model derived from our proposed j-VAR-LiNGAM method. By taking into account both instantaneous and lagged effects in the intervention, we can define instantaneous and lagged Total Effect (TE) as follows:

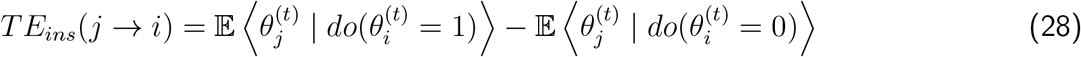

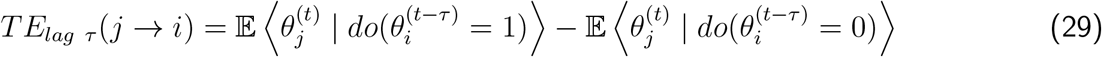

where, *TE*_*ins*_(*j* → *i*) and *TE*_*lag τ*_ (*j* → *i*) indicates the instantaneous and lagged TE from the oscillator *θ*_*j*_ to *θ*_*i*_. *τ* in Eq. (29) indicates the lagged order. Using these definitions, we can quantify the expected impact of interventions for each pair of variables in the obtained causal models. Additionally, in this study, we proposed an evaluation index called total effect centrality (TEC) to identify causally important variables within the causal model. This index is inspired by previous studies on causal inference (Runge, 2015; Ryan, 2019; Ryan & Hamaker, 2022; Ryan et al., 2018).

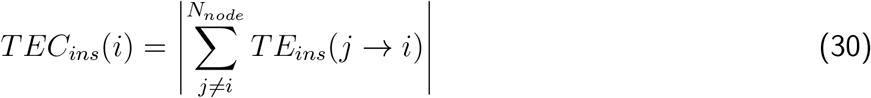

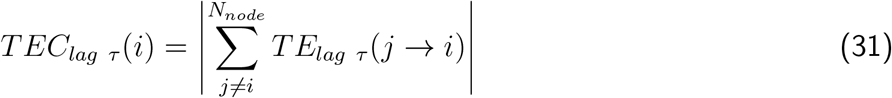

where, *TEC*_*ins*_(*i*) and *TEC*_*lag τ*_ (*i*) indicates the instantaneous and lagged TEC in the *i*-th oscillator, respectively. The increase of these values means the increase of outflow causal effect from the *i*-th to the other oscillators. Thus, we can determine the most causally important area in the causal model obtained from the j-VAR-LiNGAM, by estimating the variables associated with the maximum TEC: arg max_*i*_ : *TEC*(*i*). We presumed that the causal ordering (i.e., causal hierarchical structures) in the network dynamical systems can also be determined to refer to the ordering of TEC values. The procedures described in this section are illustrated in Fig. 1C.

To summarize our proposed approach described so far, we can establish the causal discovery and causal effect estimation method, as shown in Fig. 1A. This method can be applied to the neural data obtained from complex brain network systems since both properties, as dynamical systems and causal modeling, are rigorously considered, unlike the previous method in neuroscience studies. The Python script of j-VAR-LiNGAM we proposed is provided in the following GitHub repository: https://github.com/myGit-YokoyamaHiroshi/jVAR-LiNGAM. These scripts are required to install the lingam package (Ikeuchi et al., 2023): https://github.com/cdt15/lingam for usage.

### 2.2 Validation of the proposed method using synthetic data

To confirm whether our method can accurately extract the causal structures and causal hierarchy from network dynamical systems containing the partially bidirected feedback structures like a brain, we applied it to both numerical synthetic data and empirical neural data. In the validation with numerical simulation, we confirm whether our proposed method, j-VAR-LiNGAM, can accurately extract causal DAG structures from the synthetic observations derived from synchronous network dynamical systems, because the main aim of proposing the j-VAR-LiNGAM is to extract causally important directed information pathway from complex brain networks containing cyclic feedback structures that can be expressed as a network dynamical system. In the evaluation using empirical neural data, we applied our proposed method to neural oscillatory signals obtained from both animals and humans in vivo brains. By doing so, we confirm whether our method enables the detection of causal DAG structures and their hierarchical causal orders that reflect the interpretability of neural mechanisms.

#### 2.2.1 Simulation 1: Effect of coupling strength in synchronous network

In the first simulation, we aim to confirm the detectability of directed causal structures in our method based on the observation generated by synchronous phase-coupled oscillator models. To do this, we conduct a sensitivity analysis of our method with respect to the strength of synchronous interactions. To focus specifically on the performance of causal ordering without the influence of other components of our proposed method, we still use the j-VAR-LiNGAM-based CD method to estimate causal ordering from the observed data (i.e., steps 2 and 3);however, we omit the estimation of the dynamical model (i.e., step 1). Instead, we use the true ODE to estimate the Jacobian matrix and the steady-state solution of the phase difference that are required in step 2. Through this simulation, we primarily confirm how the estimation performance of the j-VAR-LiNGAM method affects the vicinity of the critical point in network systems among the synchronous oscillators.

##### Kuramoto-like model for generating synthetic data

The synthetic data are generated from the following Kuramoto-like oscillator model (Kuramoto, 1984; Shiozawa et al., 2022).

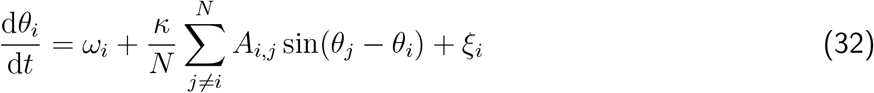

where, *θ*_*i*_, *ω*_*i*_, and *N* denote the instantaneous phase of oscillator *i*, natural frequency of oscillator *i*, and number of oscillators. *ξ*_*i*_ is observed noise drawn from a normal distribution 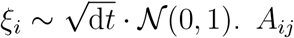 is *N* × *N* matrix that is associated with adjacency matrix of networks. The variable *κ* indicates the global coupling parameter, which controls the strength of synchronous interaction within the network systems. The critical coupling *κ*_*c*_ is given by the following equation (Shiozawa et al., 2022).

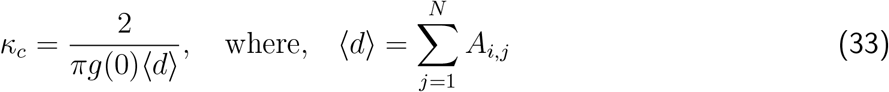

The *g*(*ω*) is standard Cauchy distribution 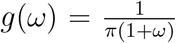, i.e., 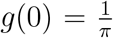. If *κ > κ*_*c*_, a phase transition from weak coupling to complete synchronization occurs among the oscillators (Kuramoto, 1984; Strogatz, 2000). In this simulation, although the adjacency matrix *A*_*ij*_ is given as fixed parameters to generate synthetic data, the parameters *κ* are changed from [0.2, 4.0] with 0.2 steps to various types of synthetic data, containing different patterns of synchronous dynamics in the networks.

##### Jacobian matrix and Fokker-Plank equation

As mentioned above, since we only focus on the performance of causal ordering, the first step of our proposed method (i.e., estimating the dynamical systems and determining their Jacobian matrices, as described in Section 2.1.1) is not performed in this simulation. Therefore, in this simulation, the true ODE (Eq. (32)) is provided as known information. The Fokker-Planck equation and Jacobian matrix are also provided as known information, derived from the Eq. (32).

The Jacobian matrix of Eq. (32) is given, below.

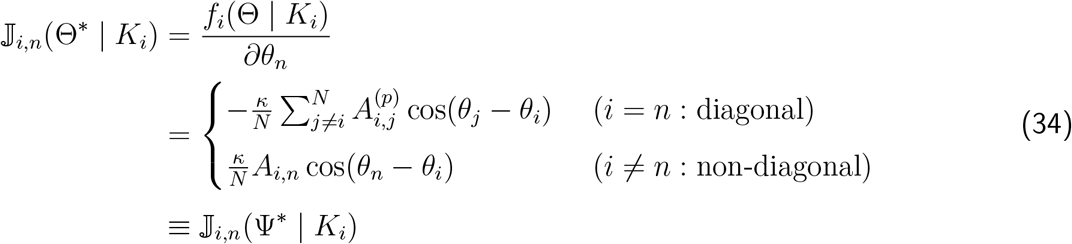

where, Θ^∗^ and Ψ^∗^ are vectors of the solution of phase and phase difference. Ψ^∗^ is obtained as numerical solution of the following Fokker Planck equation unil the distribution *p*(Ψ_*i,j*_) converges.

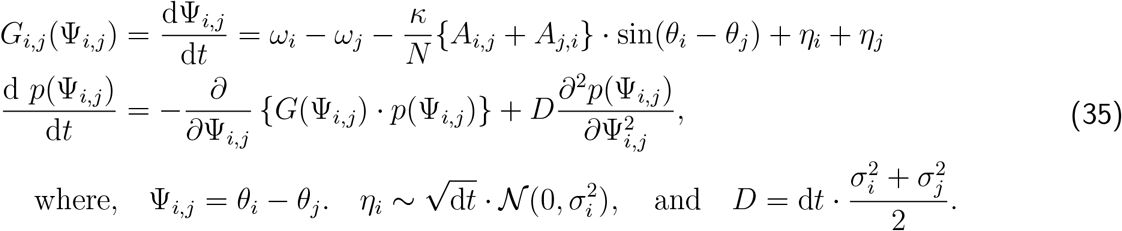

##### Simulation procedures

In this simulation, we varied the value of *κ* from the interval [0.2, 4.0] in increments of 0.2. For each value of *κ*, we generated 9,000 samples of synthetic phase data based on Equation (32) using the Euler-Maruyama method. The time step for generating the synthetic phase data was set to d*t* = 0.01. This procedure for generating synthetic data follows the same methodology as outlined in the numerical simulation described by Yokoyama and Kitajo (2022). During data generation, the natural frequency *ω*_*i*_ = 2*πf*_*i*_ of the *i*-th oscillator was fixed, with *f*_*i*_ drawn from a normal distribution 𝒩 (10, 5 × 10^−3^). The adjacency matrix *A*_*i,j*_ is set as shown in Fig. 3A (see the Result section).

**Figure 3:**
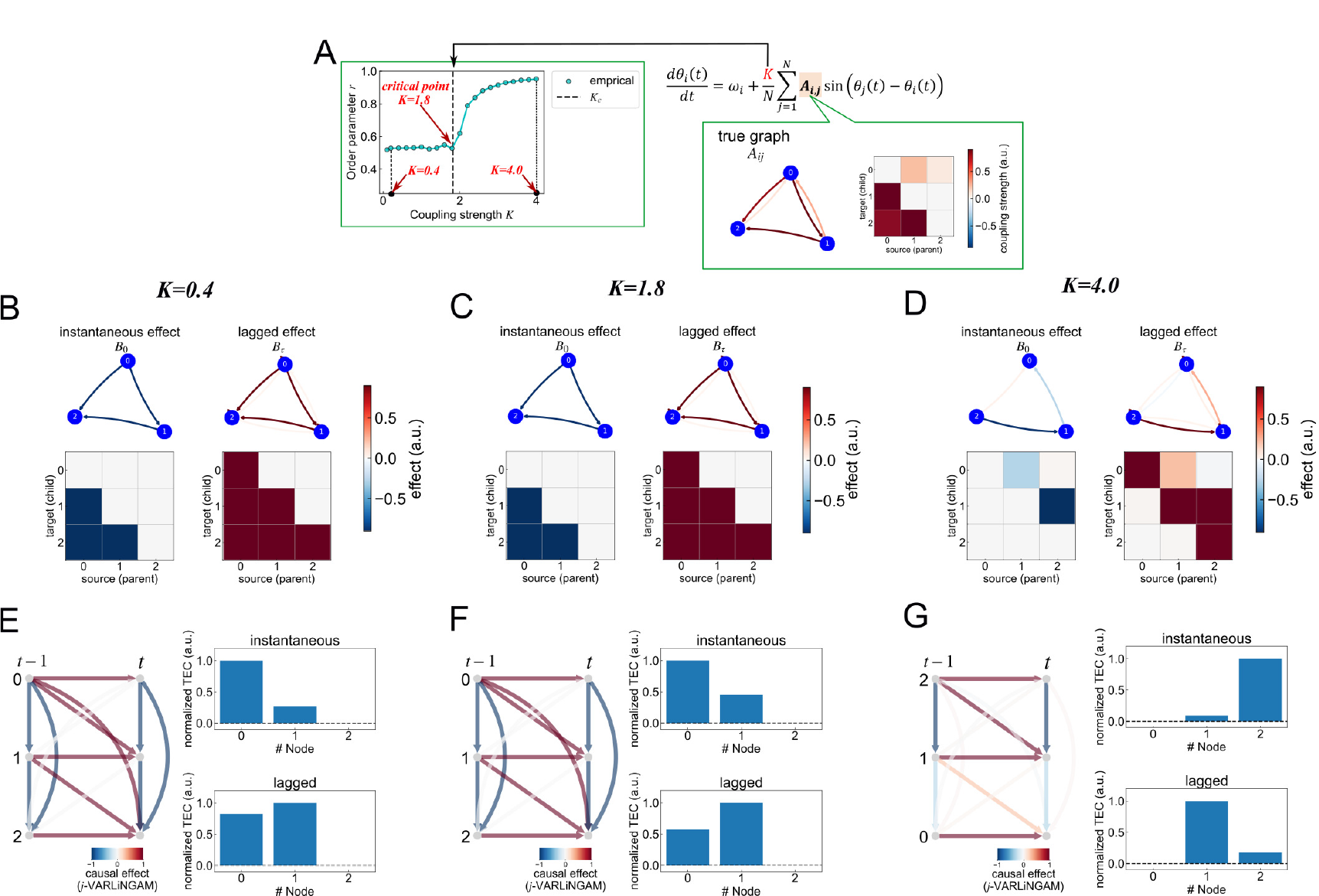
Result in Simulation 1 using j-VAR-LiNGAM. (A) True graph and phase-transition diagram in the Kuramoto-like oscillator model to generate the synthetic observations. (B–D) Estimated results in the instantaneous and lagged effect, *B*_0_ and *B*_*τ*_, for each simulation condition. The upper two panels showed the graphical diagram for each effect: *B*_0_ and *B*_*τ*_. Note autoregressive effect in *B*_*τ*_ is ignored to visualize the *B*_*τ*_. The lower two matrices showed the estimated value in *B*_0_ and *B*_*τ*_. (E–G) Time-series graph and total effect centrality (TEC). Left panel for each graph with sorting the causal ordering. The right upper and lower panels showed the instantaneous and lagged values for each normalized TEC.

After generating the synthetic phase time-series with these parameters, we applied our j-VAR-LiNGAM-based CD method to the data for each *κ*. Once the causal models were obtained using j-VAR-LiNGAM for each condition, we calculated the values of instantaneous and lagged TEC using Eqs. (28–31). This allows us to determine the variables that correspond to the causally important variable of the network for each simulation scenario.

For performance comparison, we also evaluated two baseline methods: PCMCI (Runge et al., 2023) and standard VAR-LiNGAM (Hyvärinen et al., 2010). When applying both baselines, these methods were applied directly to the synthetic data to obtain time-series causal models. Since PCMCI and VAR-LiNGAM operate on continuous time-domain data, we converted all synthetic phase time-series *θ*_*i*_ into time-domain signals *X*_*i*_ = cos(*θ*_*i*_) before applying these methods. After obtaining causal models with the baseline methods, we identified causally important variables using the same procedures and equations (Eqs. (28–31) as in our proposed approach.

#### 2.2.2 Simulation 2: performance comparisons in more complex network data

The purpose of this simulation is to test the validity of the whole procedure of j-VAR-LiNGAM-based CD method using synthetic observed data derived from synchronous network dynamical systems partially containing the cyclic connection. The synthetic data is generated from a phase-coupled oscillator model with predetermined parameters and network structure. Additionally, we will compare the performance of our method against baseline approaches (PCMCI and standard VAR-LiNGAM).

##### simulation procedures

In this simulation, we generated 9,000 samples of synthetic phase data based on the following equation, using the Euler-Maruyama method with the time-step d*t* = 0.01.

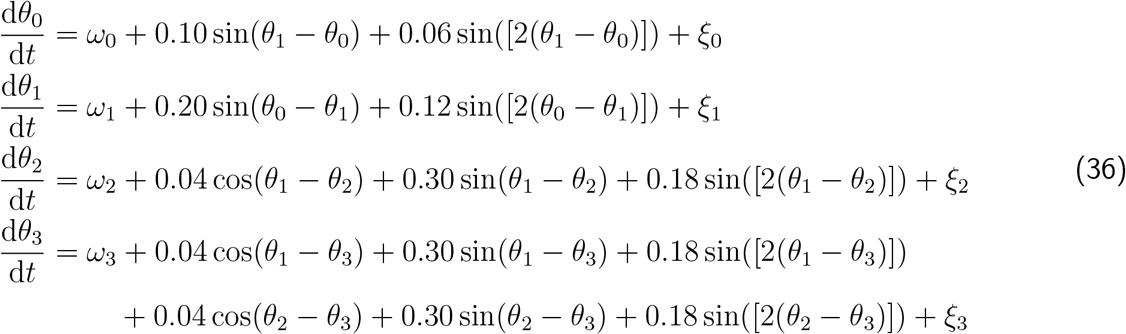

where, *ω*_*i*_ = 2*πf*_*i*_ is the natural frequency of the *i*-th oscillator with fixed value of *f* = [*f*_1_, *f*_2_, *f*_3_, *f*_4_] = [9.900, 10.505, 10.690, 10.835]. *ξ*_*i*_ is observed noise drawn from a normal distribution 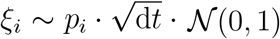, where, the noise scale *p*_*i*_ set as *p*_*i*_ = 0.05.

After generating the synthetic phase time-series with these parameters, we applied each CD method, j-VAR-LiNGAM, PCMCI, and standard VAR-LiNGAM to the synthetic observed data in the same manner in Simulation 1. After doing that, we compared the performance of causal analysis for each CD method.

### 2.3 Validation of our method using LFP data

#### 2.3.1 Aim of this validation

Next, to assess the applicability of our proposed method to high-spatiotemporal animal neurophysiological data, we analyzed laminar LFP recordings that we obtained from a mouse. Especially, using our proposed method, we analyzed each LFP data with 3-minute intervals during a foraging navigation task in a virtual reality (VR) environment and voluntary free moving in a dark room. By doing so, our objective was to test whether our method can detect hierarchical neural information pathways among the primary visual cortex (VISp), the dorsolateral geniculate nucleus (LGd), and the hippocampus (CA1, CA3) areas during the navigation task. Since the mouse is required to explore the optimal route in virtual maze on the VR environment to obtain for the reward (water), the brain in the mouse must integrate both (1) cognitive processes with sensory input and (2) the continual updating of predictions based on contextual information derived from visuomotor memory and prior knowledge in the VR environment. The former process, known as the “bottom-up process,” operates automatically, denoting the flow of information from lower-level to higher-level sensory areas (e.g., retina (visual input)→LGd→VISp→higher order visual cortex). Conversely, the latter process, termed the “top-down process,” is considered an active mechanism involving feedback modulation from higher-level to lower-level cortical areas, to minimize behavioral “prediction errors” (e.g., memory-guided visuomotor perception mediated by hippocampal (CA1/CA3) feedback to VISp). While recent studies have indicated that both bottom-up and top-down processes are present in the hierarchical flow of information between CA1/CA3 and VISp for visuomotor processing, it is unclear how both processes interact and are reflected in the functional pathway between these cortical areas during predictive behavior. Based on this issue, we applied the j-VAR-LiNGAM method to the laminar LFP signals among VISp, LGd, and CA1/C3 to estimate the hierarchical information pathway in these brain regions underlying the interaction between “bottom-up” and “top-down” processes during the task. We expected that “top-down” processes (i.e., CA1/CA3→VISp) are dominant against the “bottom-up” process (VISp→CA1/CA3) during the VR task because of the processing of the memory-guided visuomotor perception.

#### 2.3.2 Animal subject

Experiments were conducted after receiving approval from the Animal Care and Use Committee of Nagoya University. All animal procedures were conducted according to institutional and national guidelines. In this study, we used a C57BL/6J mouse purchased from Nihon SLC. Mice were housed in a temperature-controlled room (24°C ± 1°C) under a 12 h light/dark cycle with *ad libitum* access to food. Both male and female mice were used in this study (adult mice aged 20–40 weeks at the time of the imaging session). For the navigation task, water restriction is applied to the subject. During experiments, subjects were water-restricted to 85% of their free-feeding body weight. Surgery for head-plate implantation was performed as described previously in Takeuchi et al. (2025). A custom-made stainless head plate was implanted onto the skull to ensure stable head fixation under deep anesthesia with a mixture of medetomidine hydrochloride (0.75 mg/kg; Nihon Zenyaku), midazolam (4 mg/kg; Sandoz), and butorphanol tartrate (5 mg/kg; Meiji Seika). During surgery, the eyes were covered with ofloxacin ointment to prevent inadvertent injury. Body temperature was maintained by a head pad. The skull surfaces were covered with transparent dental cement, and the with white dental cement (Super-Bond, Shofu). The training phase started after 7 days post-surgery.

#### 2.3.3 Laminar LFP recording

Laminar LFP signals were collected with 2,500 samples per second using a 384-channel Neuropixels 1.0 probe connected to a PXI-based system (Jun et al., 2017). The probe was inserted 4.0 mm below the cortical surface to measure LFP signals in primary visual cortex (VISp), hippocampus (CA1, CA3), and dorsolateral geniculate nucleus (LGd), using micromanipulator. The planning of the insert trajectory was performed by pinpoint (Birman et al., 2023), and mapped to the Allen Mouse Brain Atlas (Allen Mouse Brain Common Coordinate Framework version 3.0: CCFv3; Q. Wang et al. (2020)). All data acquisition process for LFP signals was preformed with the Open-Ephys GUI. Offline filtering and further signal preprocessing for offline analysis were employed using a spike -interface Python packages.

#### 2.3.4 Experimental environment and task procedures

A custom-made virtual reality system was used for training and recording experiment, as previously described in Takeuchi et al. (2025). During the first training phase, water was automatically delivered at a reward landmark position. After mice learned to obtain the reward, it was modified to be triggered by licking at the landmark location. Rewards were granted only once per lap. In the recording session, after the mouse finished the navigation task block, neural activity and behavior during the voluntary movement in the dark environment were also recorded. See the descriptions in original papers (Takeuchi et al., 2022, 2025) for more details of task procedures and data acquisition of Laminar LFPs.

#### 2.3.5 Preprocessing and analysis

We first applied a notch filter for line frequency and its harmonics (60 and 120 Hz) to reduce the hum noise effects. After removing hum noise effect and applying anti-aliasing, EEG data were divided into epochs with 3 minute interval and transformed to current source density signals by using Python implemented script of the inverse current source density (iCDS) method proposed by (Pettersen et al., 2006) (available at https://github.com/espenhgn/iCSD). We then reduced the number of observations from 384 to 32 by averaging the LFP signals for each channel group with a region of interest (ROI) determined with Allen CCFv3. These CDS observations of LFP were resampled from 2,500 to 100 Hz for each epoch. After that, to extract theta frequency oscillations, the CSD data were band-pass filtered between 5 and 8 Hz with a zero-phase finite impulse response filter (the number of taps = 601; transition with 0.1 Hz; window function = Hanning window), and a Hilbert transform was applied to extract the time-series of instantaneous phase in the theta frequency band. The bandwidth to extract theta oscillation was selected based on the averaged peak frequency of the distribution in the power spectral density in all CDS.

After conducting the above preprocessing, the phase time-series on theta oscillations were applied to our proposed method in the same manner as Simulation 2 as described in Section 2.2.2.

### 2.4 Validation of our method using EEG data

#### 2.4.1 Aim of this validation

Finally, to assess the applicability of our proposed method to human neurophysiological data, we analyzed open-access scalp EEG recordings, which are available at Guttag (2010). We mainly analyzed 30-second intervals prior to the onset of epileptic seizures in the data from patient chb15 (file ID: 06). Seizure onset was identified through manual annotation by a clinical investigator in the original study (Guttag, 2010; Shoeb, 2009). For this analysis, we focus on oscillations in the theta band (4–8 Hz) in EEG signals, since these signals contain the most prominent ictal onset patterns in this frequency band, particularly on channels T7–P7 in the temporal region. The main aim of this validation is to test the neurophysiological applicability in humans for our proposed method by revealing whether the method correctly detects the T7-P7 channels as a causally important area for seizure-related network dynamics reflected in theta band oscillations.

#### 2.4.2 Preprocess and main analysis

EEG data containing the dataset we used were recorded with a 250 Hz sampling frequency with a 16-bit resolution (Guttag, 2010; Shoeb, 2009) using the 23 electrode aligned with the international 10-20 system bipolar montage. We first applied a notch filter for line frequency and its harmonics (60 and 120 Hz) to reduce the hum noise effects. After removing the hum noise effect and applying anti-aliasing, the EEG data were resampled from 250 to 100 Hz. To extract theta frequency oscillations, EEG data were band-pass filtered between 6 and 7 Hz with a zero-phase finite impulse response filter (the number of taps = 6,001; transition with 0.1 Hz; window function = Hanning window), and a Hilbert transform was applied to extract the time-series of instantaneous phase in the theta frequency band. All preprocessing described above is conducted by using Python programming language with the mne-Python package (Larson et al., 2025) and in-house custom scripts with scipy and numpy packages.

After conducting the above preprocess, we selected typical 15 electrodes (F7, T7, F3, C3, P3, Fz, Cz, Pz, F4, C4, P4, F8, P8, P7 and T8) in 23 phase time-series in theta EEG oscillations, and extract 30-second intervals prior to epileptic seizure onset in the data associated with the patient chb15 (file ID: 06). These phase time-series data were applied to our proposed method as same manner of Simulation 2 as described in Section 2.2.2.

## 3 Results

### 3.1 Simulation 1

As mentioned in Section 2.2.1, we first applied our proposed method to synthetic data generated by a Kuramoto-like simple phase-coupled oscillator model of three oscillators with varying global coupling parameter conditions. This simulation aimed to reveal whether our method can correctly extract causally important DAG structures from synchronous network dynamical systems like a complex brain network. Moreover, by conducting sensitivity analysis of our method for the global coupling parameter *κ*, we also show how the estimation performance of our method is affected by the phase transition dynamics around the critical point. As we described in Section 2.2.1, in this simulation, to isolate the performance of causal ordering from the influence of other components of our proposed method, Step 1 of our method is omitted from causal analysis in this simulation, i.e., the ODE equation and its parameters are given as known information to determine the Jacobian matrix and steady state of phase difference in the systems. Moreover, the performance of causal ordering estimation in our proposed method was compared with the baselines (standard VAR-LiNGAM and PCMCI). The results of this simulation are shown in Figs. 3–5.

First, we presented the results from our proposed method, j-VAR-LiNGAM (see Fig. 3). As illustrated in Fig. 3A, the synthetic observed data includes phase time-series derived from three oscillators. The true network structures in the synthetic observed data were given as shown in the true graph *A*_*i,j*_ in Fig. 3A. The synthetic data exhibit a strong directional coupling represented as 0 → 1 → 2, but also contain two weak feedback loops: 2 → 0 and 1 → 0. Moreover, by introducing the global coupling *κ*, the dynamics of the synchronous network in this model were drastically changed with increasing global coupling *κ*. Since the phase transition occurs with *κ*_*c*_ = 1.8 in this simulation, phase dynamics in the model changed from weak synchronous interaction to perfect synchrony in *κ* ≫ 1.8. If our method can accurately extract the directed acyclic causal structures without the influence of these weak feedback connections on the 0-th oscillator, it should successfully identify a DAG with the structure 0 → 1 → 2 by estimating the instantaneous effect, *B*_0_, using the j-VAR-LiNGAM approach. As shown in Fig. 3B, C, our method can correctly extract the DAG structure, 0 → 1 → 2, in the instantaneous effect, *B*_0_ (see left side graphical diagram and adjacency matrix in Fig. 3B, C). We found the tendency that the weak feedback effect for the 0-th oscillator seems to be reflected in the lagged effect *B*_*τ*_ (see the upper graphical diagram on the right-hand side in both Fig. 3B, C). Moreover, as shown in Fig. 3E, F, the causal ordering, 0 → 1 → 2, was also reflected in the ranking of instantaneous TEC. Interestingly, while the order of instantaneous TEC was consistent with the causal order, the lagged TEC would not necessarily be. In addition, when *κ* = 4.0, we found that our method cannot detect both causal DAG structure and causal order. It can be considered that the causal direction cannot be identified under the perfect synchrony condition because of the phase-locking. In this condition, three time series of the oscillators are perfectly correlated and synchronized. Thus, the inability to identify network and causal structures under perfect synchrony is a general limitation of all methods, not unique to our approach. This result showed that our method can accurately identify the causal structures in synchronous network dynamical systems under the weak coupling condition *κ* ≤ *κ*_*c*_ (*κ*_*c*_ = 1.8).

Next, to compare the performance of our method, we applied two baselines: standard VAR-LiNGAM and PCMCI to the same synthetic data. The results of standard VAR-LiNGAM were shown in Fig. 4. In this evaluation, we attempted to possibly meet the conditions for estimating the SVAR model between standard VAR-LiNGAM and j-VAR-LiNGAM, except for incorporating information from the Jacobian matrix to calculate *B*_0_ in the j-VAR-LiNGAM. Therefore, in this simulation, we set the maximum lag order for the standard VAR-LiNGAM to *τ* = 1 as well as j-VAR-LiNGAM. By doing so, we aimed to specifically reveal the effect of the causal analysis performance due to incorporate the Jacobian matrix information for calculating the *B*_0_ into the VAR-LiNGAM algorithm. As shown in the results in Fig 4B–D, regardless of the effect of phase transition, this method cannot detect causal DAG structures under any condition of the parameter *κ*. Neither the time series causal graph nor the order of TEC (both instantaneous and lagged TEC) reflected true causal structures.

**Figure 4:**
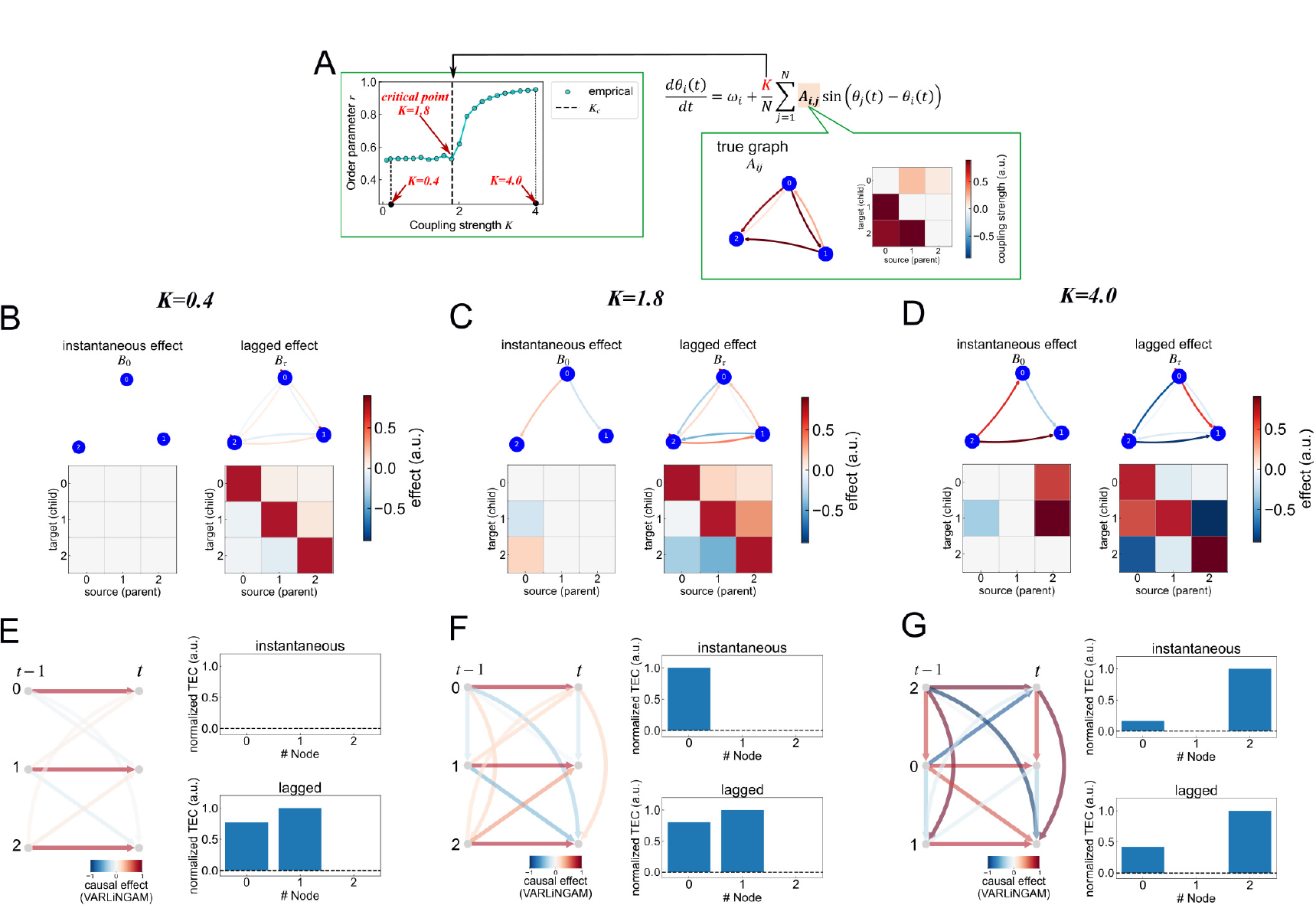
Result in Simulation 1 using baseline: VAR-LiNGAM. (A–G) Visualize the simulation results in VAR-LiNGAM in the same manner in Fig.3

On the other hand, even though the PCMCI was also not able to correctly estimate the causal structures in this simulation, a different tendency was found when applying the standard VAR-LiNGAM. Unlike our proposed j-VAR-LiNGAM and standard VAR-LiNGAM, the PCAMCI cannot uniquely detect the instantaneous effect *B*_0_; this method therefore estimates lagged causal effect based on a conditional independence test using an extended version of the PC algorithm. As shown in Fig. 5B, C, this method correctly detects similar graph structures in the true graph *A*_*i,j*_ (see Fig. 5A) as the lagged effect under the weak coupling condition *k* ≤ *κ*_*c*_ (*κ*_*c*_ = 1.8). However, as shown in Fig. 5E–F, causal order cannot be detected as 0 → 1 → 2 because the highest value of lagged TEC was found in the 1st oscillator. Therefore, PCMC can also not extract the causally important DAG structures in the dynamical systems.

**Figure 5:**
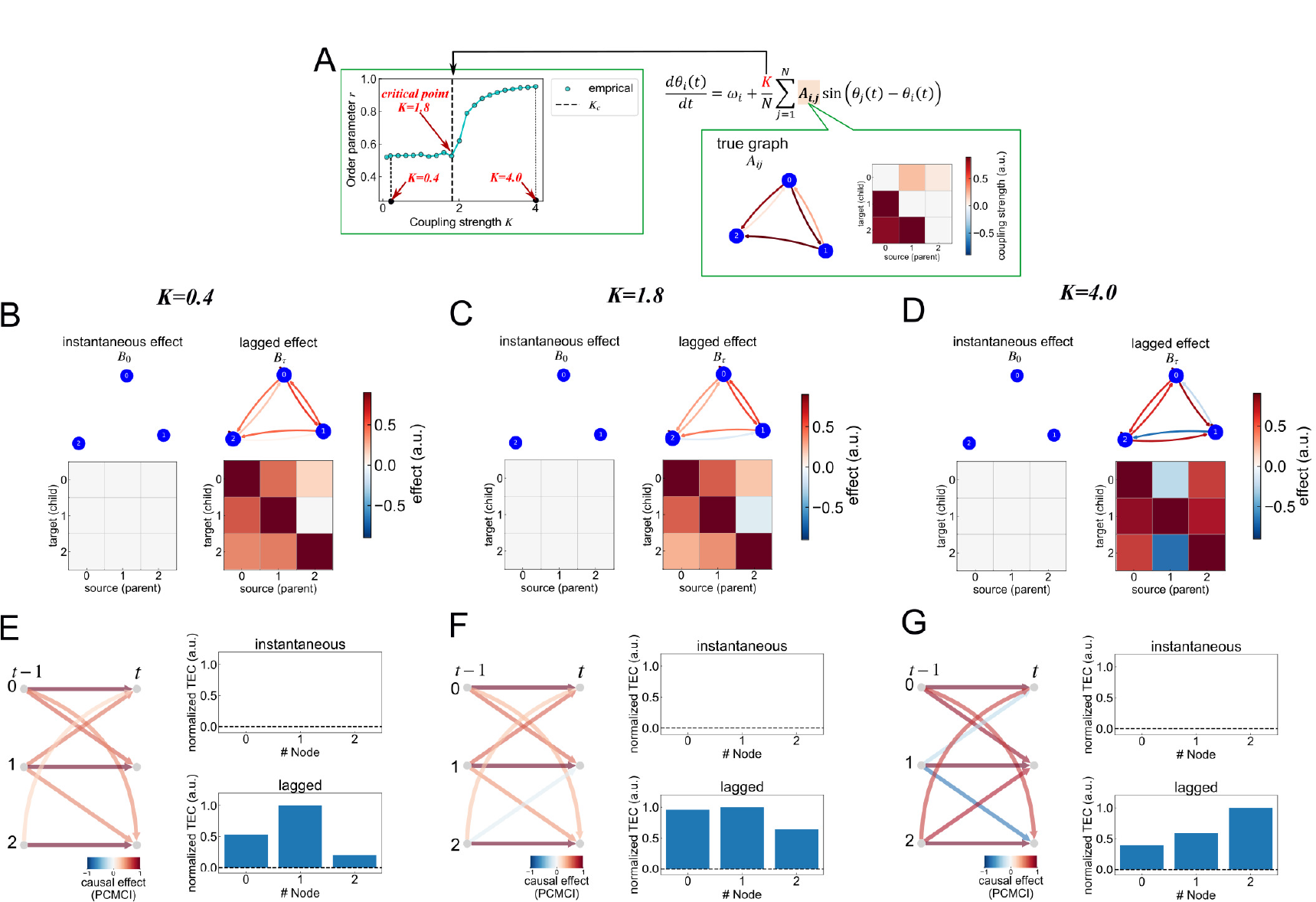
Result in Simulation 1 using baseline: PCMCI. (A–G) Visualize the simulation results in VAR-LiNGAM in the same manner in Fig.3. Note that causal orders were not sorted in Fig. 5E–C, since the topological order cannot be estimated in PCMCI.

### 3.2 Simulation 2

Our aim in this simulation is to assess the validity of the whole procedure in the j-VAR-LiNGAM method by applying it to synthetic data generated by the phase-coupled oscillator model in Eq. (36). As in Simulation 1, we tested whether our method accurately extracts causally important DAG structures from network dynamical systems with partial cyclic feedback. The synthetic data partially included directed connections as shown in Fig. 6A (see the link between the 0th and 1st oscillators). If our method works as intended, it will detect the correct causal order: 0 → 1 → 2 → 3, unaffected by feedback from the 1st to the 0th oscillator.

**Figure 6:**
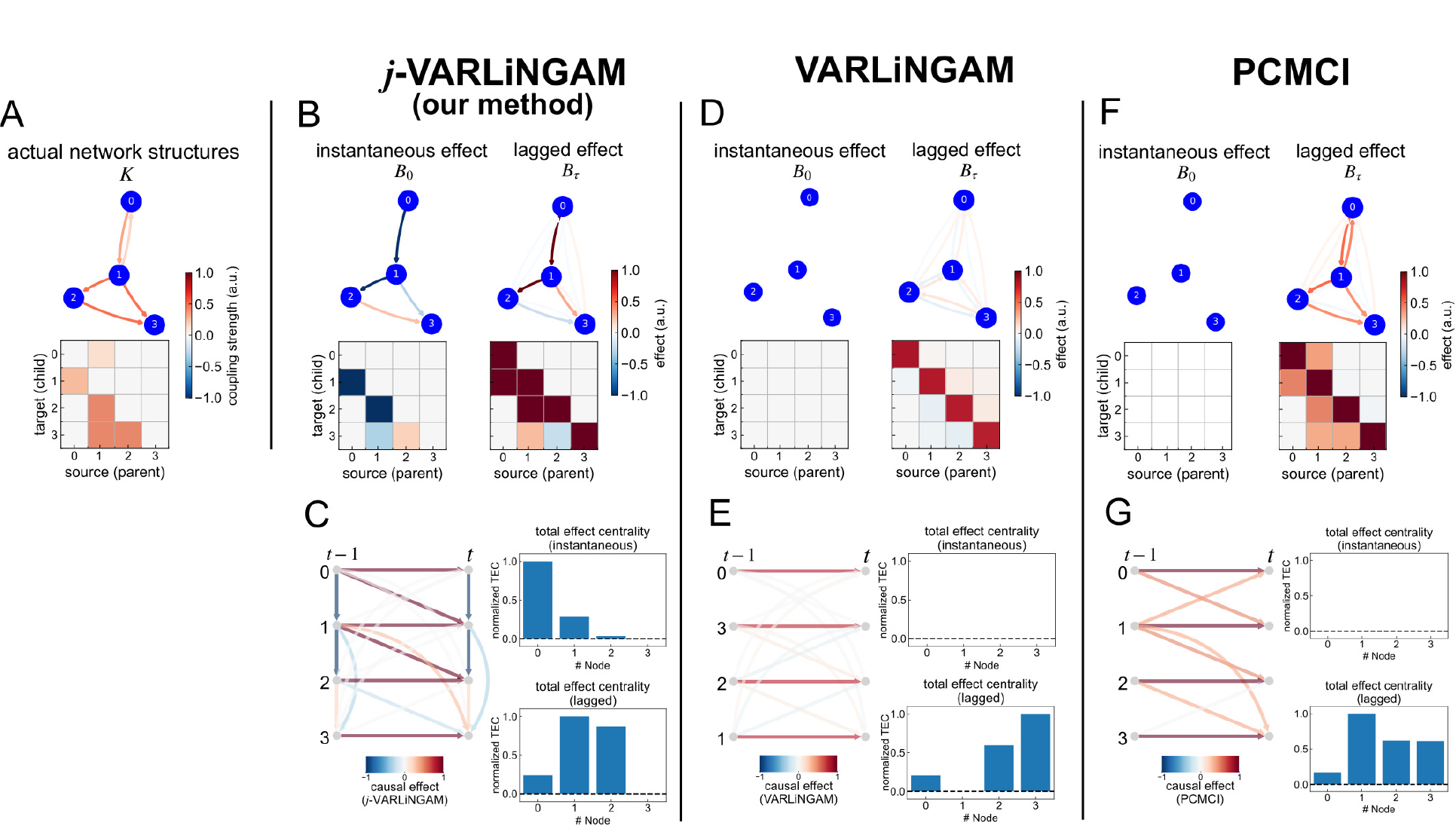
Results in the Simulation 2. (A) True graph and phase-transition diagram in Kuramoto-like oscillator model to generate the synthetic observations. (B–D) Estimated results in the instantaneous and lagged effect, *B*_0_ and *B*_*τ*_ for each simulation condition. Upper two panel showed the graphical diagram for each effect: *B*_0_ and *B*_*τ*_. Note autoregressive effect in *B*_*τ*_ is ignored to visualize the *B*_*τ*_. Lower two matrix showed estimated value in *B*_0_ and *B*_*τ*_. (E–G) Time-series graph and total effect centrality (TEC). Left panel for each graph with sorting the causal ordering. Right upper and lower panels showed the instantaneous and lagged value for each normalized TEC.

As a result, our j-VAR-LiNGAM method successfully identified the intended causal DAG. As shown in the left panel of Fig. 6B, the instantaneous effect *B*_0_ accurately captured the causal order: 0 → 1 → 2 → 3. Moreover, the descending order of the instantaneous TEC value accorded with the causal order (Fig. 6C), as in the result of Simulation 1. In contrast, as shown in Fig. 6 D–G, while our method consistently determined the correct causal order from observed data, the baseline methods (standard VAR-LiNGAM and PCMCI) failed to extract both the DAG structure and the correct ordering. In particular, as shown in the right panel of Fig.6F, PCMCI determined similar network structures to actual network structures (Fig.6A); however, the causal order cannot be determined as we intended from estimation results in lagged TEC (see bottom of right panel in Fig.6G).

In summary of both results in Simulation 1 and Simulation 2, these results suggested that our method, j-VAR-LiNGAM, can accurately extract the causally important DAG structures from the synchronous dynamical systems with partial cyclic feedback, as in the complex brain network.

### 3.3 Mouse Laminar LFP

To confirm the neurophysiological validity of the j-VAR-LiNGAM method, we then applied this method to the laminar LFP data we recorded from an in-vivo mouse brain. As outlined in Section 2.3, this validation aims to assess the applicability of neurophysiological data by revealing whether our method detects the hierarchical information flow reflected in the local circuit among VISp, CA1/CA3, and LGd. Specifically, we focused on information flow mediated by memory-guided visuomotor perception. The j-VAR-LiNGAM method was employed on LFP data recorded during both the foraging navigation task in a virtual reality (VR) environment (i.e., memory-guided visuomotor task) and the voluntary free-moving task. By comparing the results, we attempt to clarify the causal structures in the local circuit associated with this perception.

The results in the VR foraging navigation task were shown in Fig. 7. Fig. 7B, C illustrates the estimated time-series causal graph and TEC analysis obtained by using the j-VAR-LiNGAM method. For visualizing the time-series causal graph in Fig. 7B, the order of ROI in the observed data was aligned with the causal ordering of the causal graph estimated by our method. As can be seen in both results in Fig. 7B, C, the ROIs associated with CA1 (ROI 19), LGd (ROI 5), and VISp (ROI 25) were determined as higher-order areas in the causal hierarchy. We found that the ROI 19 related to the CA1 showed the largest value in the instantaneous TEC and the highest order of the hierarchy. To further examine the significant flow of causal information from these ROIs, we visualized a topological sorting graph derived from the resulting causal Directed Acyclic Graph (DAG) in our analysis, as shown in Fig. 7C. For this visualization, we displayed only those causal edges with a larger effect that satisfy the condition 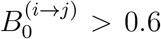. We specifically highlighted the causal pathway originating from the regions that include CA1 (ROI 19), LGd (ROI 5), and VISp (ROI 25). See the red bold directed edges in Fig. 7C.

**Figure 7:**
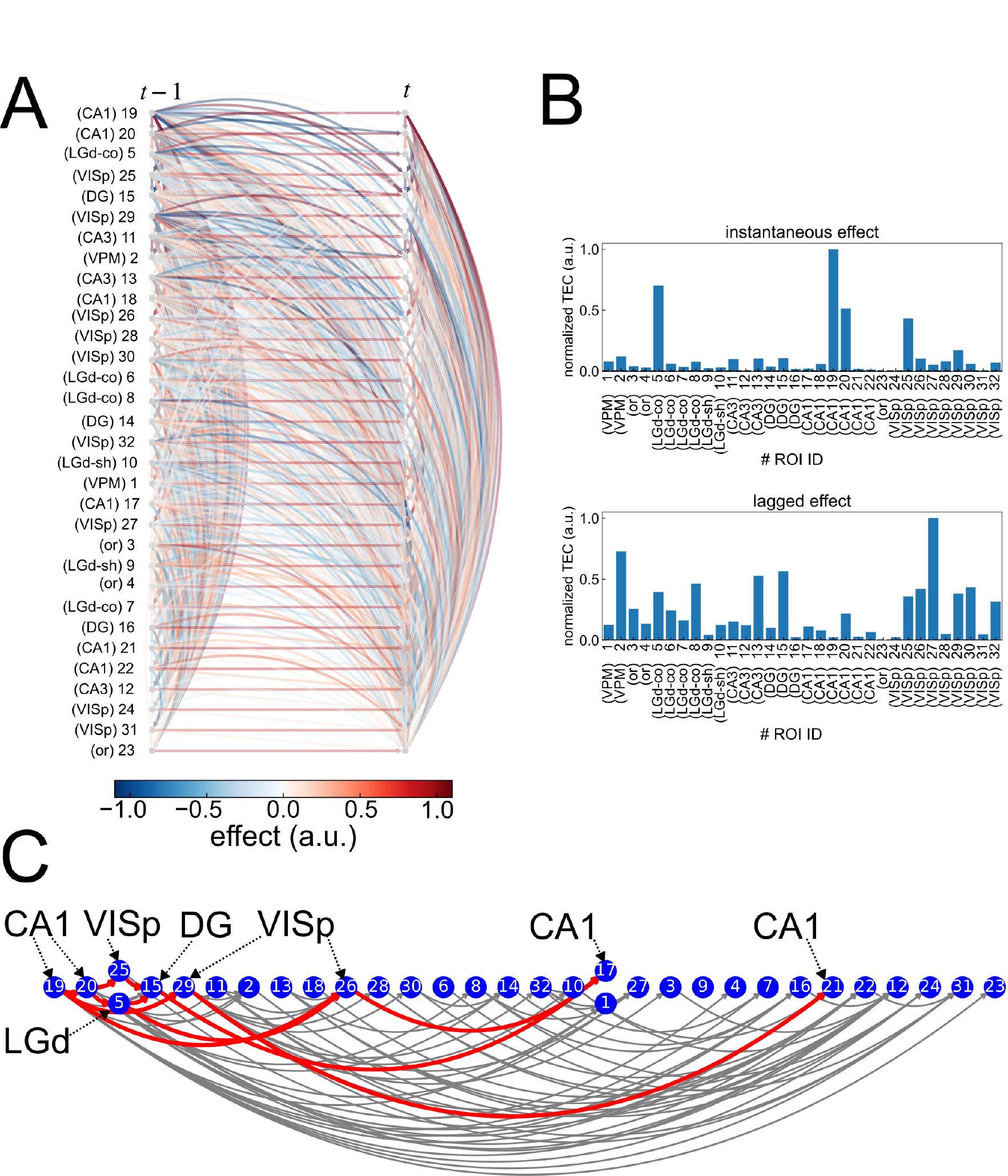
Causal analysis in the mouse laminar LFP during VR foraging navigation task. (A) Time-series causal graph obtained by the j-VAR-LiNGAM. The order of nodes in the graph is sorted according to their causal order. (B) Instantaneous and lagged TEC results. The value for each TEC is normalized with the maximum value of TEC. (C) Topological sorting of the graph in the mouse laminar LFP. The red and gray colored edges of this graph illustrate the causal pathway containing the higher causal effect conditioned by 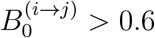. The red highlighted edges indicate the specific causal pathway originating from CA1 (ROI 19) that is the hierarchically most upper ancestor in the causal graph. The abbreviation of ROI labels in (A–C) indicates as follows: VISp: primary visual cortex, CA1, CA3: the first and third region in the hippocampal circuit, DG: dentate gyrus, LGd: dorsolateral geniculate nucleus, and VMP: ventral midbrain*/*pons.

As shown in Fig.7C, these highlighted edges in the topological sorting graph indicated the following two key findings. First, the first five ROIs in this graph (containing ROI IDs: 19, 20, 5, 25, and 15) were organized to be modular structures, indicating the hippocampal (CA1) feedback to the VISp (i.e., indicating the top-down processing). Second, the ROI 19 (CA1) has the causal pathway: CA1(ROI:19)→VISp(ROI:29)→CA1(ROI:21) and CA1(ROI:19)→VISp(ROI:26)→CA1(ROI:17). Consequently, these two connections indicates the possible roles in hippocampal top-down connection to VISp (i.e., 19→29, 19→26) and its feedback effect from VISp to CA1 (i.e., 29→21, 26→17). This result can be considered as the top-down processing modulating the bottom-up processing by propagating visuomotor memory-guided feedback.

In contrast of the results in the VR foraging navigation task, we found a quite different tendency in the voluntary free moving task. The results in the voluntary free moving task were shown in Fig. 8. As can be seen in Fig. 8A, the highest ancestor node in obtained time-series causal graph in this task were selected as the ROI 7 associated with LGd. In the TEC analysis, we also found that ROI 7 showed the largest value in the instantaneous TEC. Interestingly, unlike the results in Fig.7B, Fig.8B showed that the hierarchical dominance of ROI associated with CA1 disappeared during voluntary free moving, because the value of instantaneous TEC in ROI 7 is quite larger than other ares. To further reveal such difference in the results in VR task, we then confirmed the topological sorting graph in the voluntary free moving task (Fig.8C). This topological sorting graph suggested the following key findings. Fig.8C illustrated the topological sorting graph which was highlighted the causally significant pathway originating from LGd (ROI 7). See the bold red edges in Fig.8C. As in this figure, we found two causally significant path from ROI 7 (LGd): 17(LGd)→25(VISp), and (2) 17(LGd)→16(DG)→19(CA1). Since the LGd plays a role in mediator to propagate the visual input from retina to VISp, the first connection: 17(LGd)→25(VISp) can be considered the typical visual processing related bottom-up pathway. The later one: 17(LGd)→16(DG:dentate gyrus)→19(CA1) can be also considered as the functional bottom-up connections to send the visual perception to hippocampus. The DG is a subfield of the hippocampal formation that plays a role in the primary input region for the hippocampus. Moreover, as we mentioned in Section 2.3, LGd mediates to propagate the visual perceptual input from retina to visual cortex. Namely, the connections from LGd to CA1 mediated by DG reflected in the causal path: 17(LGd)→16(DG)→19(CA1) is reasonable from physiological perspective in visual perception processing and visual perception related bottom-up processing among these ROIs, despite of no direct anatomical connection between LGd and DG.

**Figure 8:**
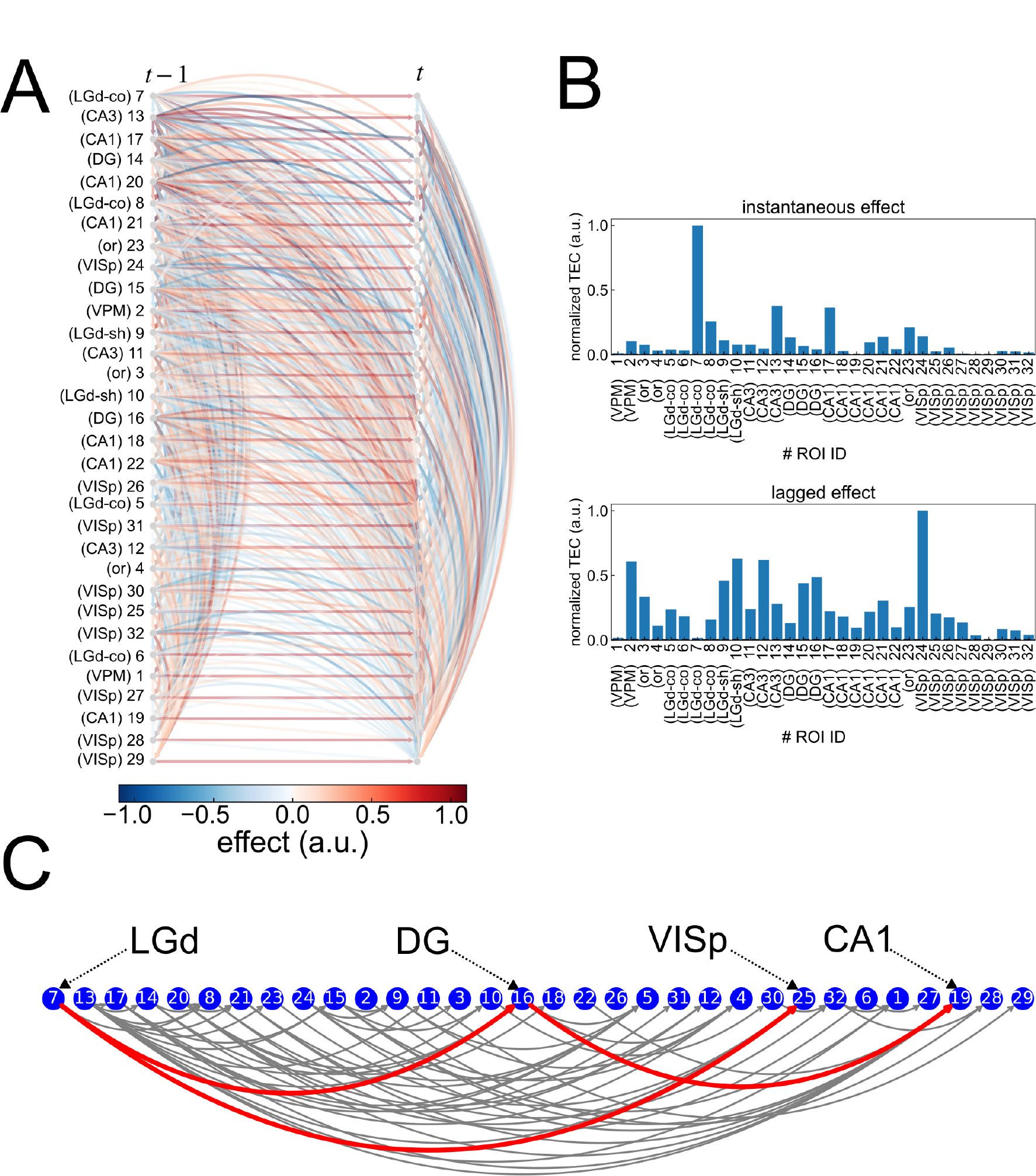
Causal analysis in the mouse laminar LFP during voluntary free-moving task. (A) Time-series causal graph obtained by the j-VAR-LiNGAM. (B) Instantaneous and lagged TEC results. The value for each TEC is normalized with the maximum value of TEC. (C) Topological sorting of the graph in the mouse laminar LFP. The results in (A–C) are visualized in the same manner in Fig.7. The red highlighted edges in (C) indicate the specific causal pathway originating from LGd (ROI 7) that is the hierarchically most upper ancestor in the causal graph.

### 3.4 Human scalp EEG

Finally, we provided the analysis results in the human scalp EEG before the epileptic seizure onset using the j-VAR-LiNGAM method. The estimated causal graph structures are shown in Fig.9A, B. While the time-series causal graph was illustrated in Fig.9B, the causal graph structures visualized on an actual montage of the EEG electrodes were shown in Fig.9A. The causal graph with EEG montage in Fig.9A separately illustrates the instantaneous and lagged effect, *B*_0_ and *B*_*τ*_ (note: the autoregressive effect in the lagged effect *B*_*τ*_ was ignored to visualize in this figure, see the right panel of Fig.9A). As described in the Section 2.4, the most prominent ictal onset patterns in the dataset we used were reflected in the theta frequency band oscillation in the T7–P7 area. Therefore, as in the qualitative trend, the larger effect causal path from T7 or P7 can be found in Fig.9A, B. As a result, the largest values of instantaneous TEC were associated with T7 electrodes. Since both results in Simulation 1 and 2 suggested that the descending order of the instantaneous TEC value accords with the causal order estimated by the j-VAR-LiNGAM method, this result suggested that the T7 electrodes are the most causally important area.

**Figure 9:**
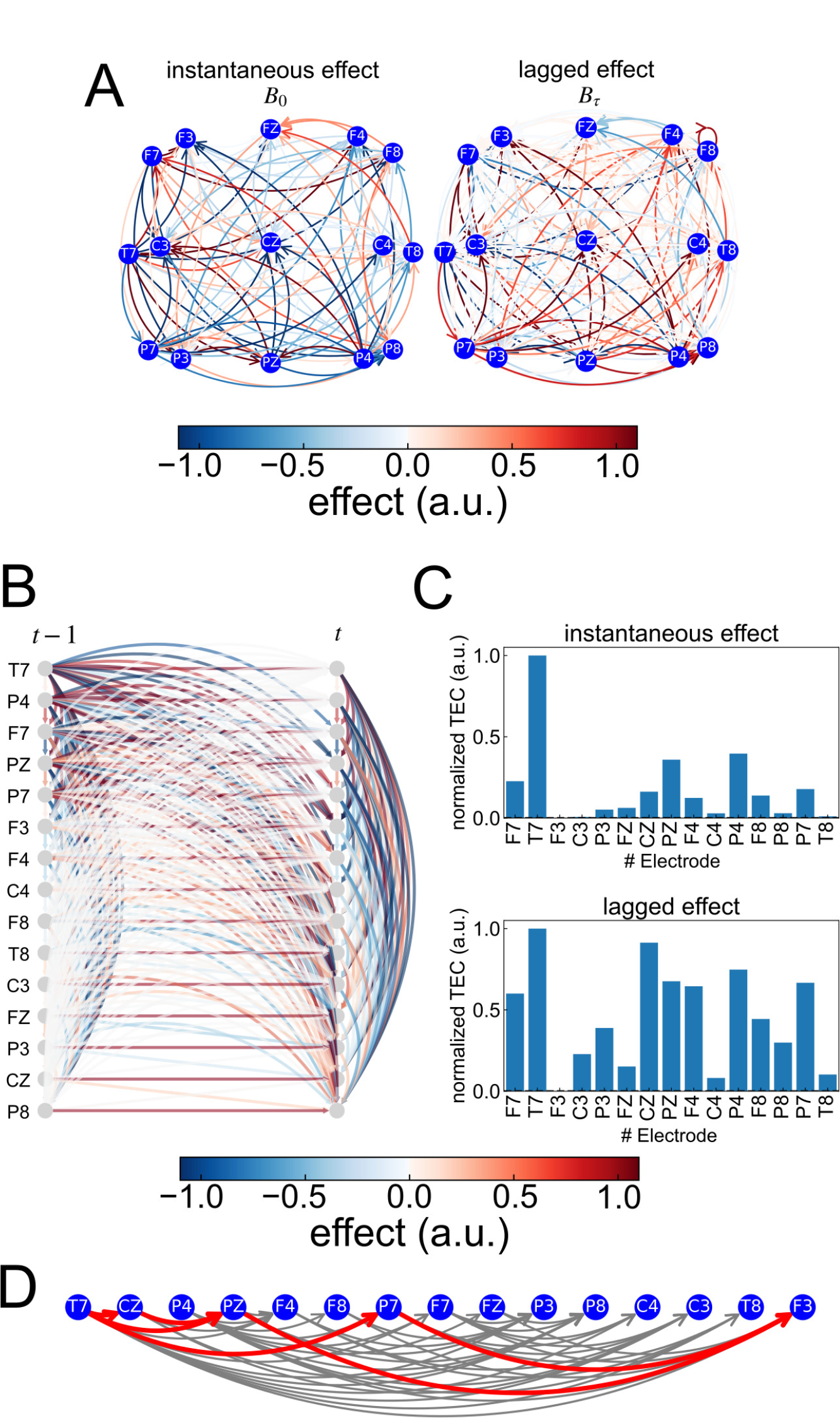
Causal analysis in the human scalp EEG before onset of the epileptic seizure. (A) Instantaneous and lagged causal graph, *B*_0_ and *B*_*τ*_, visualized on an actual montage of the EEG electrodes. (B) Time-series causal graph obtained by the j-VAR-LiNGAM. Red and Blue graduated color edges in (A, B) indicate the positive and negative direct causal effect. Note that the autoregressive effect in the lagged effect *B*_*τ*_ in (A) is ignored for the visualization. (C) Instantaneous and lagged TEC results. The value for each TEC is normalized with the maximum value of TEC. (D) Topological sorting of the graph in the EEG. This graph is visualized in the same manner in Figs.7C and 8D. The red-highlighted edges in this figure (D) indicate the specific causal pathway originating from the T7 electrode that is the hierarchically most upper ancestor in the causal graph.

To quantitatively deepen understanding of the causal information pathway from T7, we visualized the topological sorting graph of the estimated causal model using j-VAR-LiNGAM as shown in Fig.9D. For visualization, this topological sorting graph was illustrated with a causal edge of a larger causal effect to satisfy 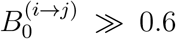. As a result, the sorted graph revealed two key findings: (1) T7 is the highest variable in the causal hierarchy of the estimated causal model, and (2) both T7 and P7, which are related to the epileptogenic zone, play a role in propagating the effects from this zone. These findings align with the trends observed in our results in Fig. 9 A–C, indicating that the j-VAR-LiNGAM method accurately identifies the T7 electrodes as a causally important area within the network (i.e., the epileptogenic zone). Additionally, it is important to note that these results were derived from analyzing EEG data collected during 30-second intervals preceding the ictal onset. Therefore, the results in this EEG analysis demonstrated the identifiability of the epileptogenic symptom before the ictal onset using the j-VAR-LiNGAM method.

## 4 Discussion

In this study, we proposed a new approach to extract causal directed network structures from observed neural oscillatory signals by combining data-driven dynamical modeling (Yokoyama & Kitajo, 2022) and the time-series CD method (Hyvärinen et al., 2010; Shimizu, 2014). The numerical validity of our proposed method was confirmed by showing that the method can accurately extract causal DAG structure and causal ordering in this DAG under the weakly coupled limit cycle states in the network dynamical systems. Moreover, the results in applying our method to animal and human neural oscillations suggested that our method enables the detection of the physiologically valid causally hierarchical structures. These findings suggest that our proposed method has theoretical and neurophysiological validity. In the following section, we will discuss the neuroscientific significance of our proposed method, and will compare our results with those of previous studies that have employed causal analysis in the complex brain network dynamics.

### 4.1 Advantages of our proposed method in neuroscience

The most significant and originality of our proposed method is that the method considers both things: (1) time-evolving dynamics reflected in the synchronous neural oscillations, and (2) causal directed structures and their identifiability. As we described in the Introduction section, even though the complex brain network contains bidirectional and cyclic interaction within the inter-cortical feedback loops, some experimental studies suggested that structural information flow in the brain is inherently directional, organized by modular and hierarchical poly-synaptic signal propagation (Greaves et al., 2025; Park & Friston, 2013; Seguin et al., 2023). This indicates that the causal directed pathway in the brain networks plays a role in the task-related or state-dependent information processing; however, the conventional method of causal analysis in neuroscience study cannot uniquely identify such directionality in the brain network because the most studies regarding causality in the brain network are based on the “GC”-based (Astolfi et al., 2008; Gourévitch & Eggermont, 2007; Granger, 1980; Schreiber, 2000) or “DCM”-based (Penny et al., 2004) perspective. As we mentioned in the Introduction section, this method does not guarantee the identifiability and directionality of causal structures in the estimated networks. Moreover, non-identifiability in these conventional methods also indicates a difficulty for estimating the causal effect in the brain network. This leads to the technical difficulty of proving the proof of concept regarding the structural information flow in the brain based on the observational and experimental neural data.

In contrast to the conventional methods in neuroscience, our proposed CD methods in network dynamical systems is identifiable under the condition satisfying the causal assumptions: (1) synchronous neural oscillations could be expressed as a ODE as like a phase-coupled oscillator model, (2) no unobserved confounders in the observations, and (3) ODE in the model can be linearized as SVAR model with Padé approximation under the non-Gaussianity in the noise. As we mentioned in the Introduction section, in the linear systems under the non-Gaussianity in the noise, the causal model is identifiable (Shimizu, 2014; Shimizu et al., 2006), and it is theoretically guaranteed because the causal model estimated under such assumptions meets the requirements of the identifiable functional model class (IFMOC) (Peters et al., 2011). Therefore, if we can consider that the causal mechanisms in the observed neural oscillations satisfy the above three assumptions of our proposed method, we can estimate the causal model with IFMOC using our proposed method, j-VAR-LiNGAM. Moreover, as we demonstrated in the numerical simulation results, we confirmed the validity and accurate detection of causal DAG structures reflected in the observations generated by the phase-coupled oscillator models under the weakly coupled limit cycle state (Fig.3). Consequently, our proposed approach, j-VAR-LiNGAM, would be identifiable to detect the causal structures in the complex brain network under the above assumptions.

### 4.2 Neurophysiological interpretability

Our proposed method advances the above-mentioned conventional methods in the following ways. Our method detects both the time-series causal structures and causally significant nodes in brain networks. It also considers the time-evolving dynamics in the brain network, which are reflected in synchronous neural oscillations. By combining two methods: the data-driven dynamical network modeling (Yokoyama & Kitajo, 2022) and the time-series CD method (Hyvärinen et al., 2010; Shimizu, 2014), we established a new approach. This method detects the causal DAG structures reflected in the steady-state of network dynamical systems formulated as phase-coupled oscillators. As a result, we can extract causal DAGs from network dynamical systems with cyclic loops and also identify interventional causal effects. The advantage of our method lies in detecting causally significant nodes in the detected time-series DAGs based on the TEC analysis we conducted.

As shown in the results in both animal laminar LFP and human scalp EEG data, the causal analysis results using our method showed the causally hierarchical information flow reflected in the synchronous neural oscillations that can make neurophysiologically valid and reasonable interpretations. In the LFP analysis, we demonstrated the hierarchical interactions that the visuomotor memory guided top-down processing (CA1→VISp) modulates the bottom-up visual sensory processing (VISp→CA1) as shown in Fig.7C. These causal pathways for visual memory guided processing obtained by using j-VAR-LiNGAM agreed with the experimental evidence. The top-down process from CA1 to VISp was reported in Fournier et al. (2020), where neural activity in the VISp is modulated by theta oscillation in CA1. The interactions between top-down and bottom-up causal pathways between VISp and CA1 regions are also suggested in Haggerty and Ji (2015), which is related to the encoding process for the spatial memory trace in the cortical-hippocampal assemblies overlapping VISp location-responsive cells and hippocampal place cells. On the other hand, during a voluntary free-moving task without VR visual feedback in a dark room, such modulation by the top-down process against the bottom-up process is not exhibited, or rather, the causal structures changed towards the bottom-up process are more dominant. As can be seen in Fig.8C, two specific causal pathways from LGd (ROI ID: 7) associated with the most causally highest ancestor are exhibited during the voluntary free-moving states: LGd→VISp (7→25) and LGd→DS→CA1 (7→16→25). The first pathway can be considered as the typical connections to process the input of visual perception from retina, since it is known as the visual perception from retina sent to the VISp area mediated by LGd (Bienkowski et al., 2019). Moreover, an experimental study suggested the connectivity between LGd–VISp is modulated by locomotion behavior (Aydın et al., 2018). Therefore, considering this experimental evidence, it is physiologically reasonable that the causal pathway in LGd→VISp is found during a voluntary free-moving task. Moreover, the causal pathway in LGd→DS→CA1 is also consistent with the experimental evidence reported in Gabriel et al. (1996). Even though this study does not directly suggest the contribution to the visuomotor processing, this study suggested that the interaction among LGd, DG, and CA3 mediated by cholecystokinin and glutamate agonists would be involved in the maintenance of functional plasticity in CA1 (Gabriel et al., 1996). Therefore, the causal pathway in LGd→DS→CA1 we found is reflected in such functional mechanisms among these regions.

Neurophysiologically interpretable results were also found in our analysis with an open data set of seizure EEG signals. In this EEG analysis, our method detected the seizure-related directed causal pathway from the epileptogenic zone, as suggested in the original paper of our used dataset (Shoeb, 2009) (see Fig.9D). Moreover, as an important finding, we can detect the electrodes associated with the epileptogenic zone as the most causally significant area in the causal graph obtained by our proposed method. This is notable because we found such a seizure-related causal pathway from the EEG theta oscillations before the ictal onset time. These results support that our method is detectable for the neurophysiologically valid causality reflected in the EEGs in epileptic patients before the occurrence of the epileptic-seizure-related theta oscillations. Moreover, as can be seen in Fig.9D, we found the seizure propagation pathway from the T7 area (i.e., the epileptogenic zone), such as in T7→PZ→F3, towards the connection from the temporal to the frontal cortex. Some recent studies on the temporal lobe epilepsy using stereo-EEG suggested the temporo-frontal connection that plays a key role in the seizure propagation pathway(Bartolomei et al., 2025; Oane et al., 2025).

In summary, by adopting the property of time-evolving dynamics in the observed data into the time-series CD method using our proposed method, j-VAR-LiNGAM, the proposed method succeeded in providing a quantitative interpretation of the causal hierarchy of time-evolving dynamics in complex brain networks in both animal and human neural oscillatory data. Therefore, our findings from both numerical simulations and neurophysiological validations indicate that our proposed method offers a promising approach for demonstrating directed information flow driven by task-related or state-dependent core brain regions, as mentioned in previous experimental studies (Greaves et al., 2025; Park & Friston, 2013; Seguin et al., 2023).

### 4.3 Significance of the proposed method in the context of causal discovery

To our knowledge, while some studies on statistical causal inference have discussed dynamical systems as causal models and their identifiability (Bongers et al., 2022; Mooij et al., 2013; Y. Wang et al., 2024), no CD method has been established to detect causal models in a data-driven manner for estimating causal structures in network dynamical systems like the brain. As noted in the Introduction, CD methods such as PCMCI were applied to nonlinear observations. However, it is unclear whether these methods are valid for synchronous network dynamics in neural oscillations, including LFP and EEG signals.

In contrast, our simulations show accurate detection and numerical validity for causally significant nodes in the graph by using our proposed method, j-VAR-LiNGAM (see Figs. 3–6). The j-VAR-LiNGAM method can detect the causal hierarchical order among observational data, even when weak feedback loops exist. This is possible even in hard detection cases for standard VAR-LiNGAM and PCMCI. This contrast highlights our method’s advantage. False detections by standard VAR-LiNGAM show the challenge of finding causal structures through direct linearization of neural data. PCMCI, while developed for model-free nonlinear time-series CD in dynamical systems (Runge et al., 2019, 2023), still faces difficulty in extracting causal structures. Thus, bringing Jacobian-matrix information into VAR-LiNGAM is crucial for extracting hierarchical causal structures from neural data in dynamical brain networks. These findings demonstrated a clear advantage of our proposed method, j-VAR-LiNGAM.

### 4.4 Limitations of this study

Despite the advantages of our method, some limitations exist. First, although we demonstrated the advantage of our method, the validity of the estimation in our method is guaranteed only when the aforementioned assumptions are satisfied. Moreover, our proposed method consists of the three steps: (1) dynamical systems identification (Yokoyama & Kitajo, 2022), (2) ODE linearization and extraction of the causal structures in the systems, and (3) causal effect estimation. Therefore, the accuracy of CD in our methods would be affected by the estimation error for each step. From a practical perspective, a large number of observation samples were required to minimize the prediction error for each step. As we priory discussed in the Supplementary Information of Yokoyama and Kitajo (2022), the dynamical systems identification in the above step (1) requires at least 4,000 times iteration for model updating when applying this method to the observations with over 30 numbers in the oscillators, because the method in the step (1) is based on the Bayesian inference as described in the Section.2.1.1. Based on these issues, we limited to apply our proposed method, j-VAR-LiNGAM, for the observations containing around 30 oscillators. Further studies are required to confirm the identifiability and estimation accuracy for applying the j-VAR-LiNGAM to the high-dimensional observations.

Certainly, we obtained the neurophysiologically valid results using our proposed method in both animal and human data. However, to enable us to establish our findings, obtained from j-VAR-LiNGAM, as”actual” neurophysiologically valid causality, further interventional studies using animals are crucial. These studies would help confirm whether the estimated regions are truly associated with causally important areas for controlling network systems in the brain. To achieve this, neurophysiological experiments need to be conducted using animals to assess the interventional effects of modulating neural activity in regions identified as causally important through the causal analysis of j-VAR-LiNGAM. Neuromodulation techniques can be practically useful for confirming the causal roles of specific brain regions. Although this aspect is beyond the scope of the current study, such validation will be essential to substantiate the “actual” neurophysiological validity of our proposed CD method in future research.

Even with these limitations, our method provides advantages over the conventional methods in that it estimates both causally hierarchical structures and causally significant nodes in the complex brain networks. To reveal the estimation accuracy in the above step (1), we confirmed that the stationary distribution of the probability *p*(Ψ) in phase difference was well fitted to the empirical distribution of phase difference obtained from the histogram of phase difference in the observational phase in oscillators. In the Supplementary Materials, we further conducted the comparisons between the observational and estimated stationary density *p*(Ψ) derived from the Fokker-Planck equation (Eq. 4). This evaluation, conducted for Simulations 1 and 2 in Sections 2.2.1 and 2.2.2, showed that the estimated *p*(Ψ) fits the empirical distributions well, confirming the accuracy of the estimation in the current study (See Figs. S1 and S2 in the Supplementary Materials).

## 5 Conclusions

In this study, we proposed a new CD method, j-VAR-LiNGAM, which can be applied to estimate the causal directed information flow in a complex brain network, while considering both dynamical properties and causal directed structures reflected in the observed neural oscillations. We confirmed the validity of our method by showing that the method accurately estimates the causal structures in a data-driven manner in both numerical and empirical data analysis. For future work, we will conduct the study on animal neuroscience to confirm the interventional effect for the regions related to the causally significant nodes in the complex brain network systems determined using our proposed method. To attempt such future studies, further consideration is required to address the above-mentioned limitations of this study.

## Data and Code Availability

The programming code to reconstruct the results in all numerical simulations and open EEG dataset was implemented using the Python language. These are available at Github: https://github.com/myGit-YokoyamaHiroshi/jVAR-LiNGAM. The raw data of mouse LFPs and the programming code for the analysis associated with these LFPs will be made available to all interested researchers upon request.

## Author Contributions

**Hiroshi Yokoyama**: Conceptualization, Data curation, Formal analysis, Funding acquisition, Investigation, Methodology, Resources, Software, Validation, Visualization, Writing original draft, Writing – review & editing. **Ryosuke Takeuchi**: Data curation, Investigation, Methodology, Writing – review & editing. **Shohei Shimizu**: Project administration, Supervision, Resources, Software, Validation, Writing – review & editing.

## Acknowledgements

This research was partially funded by the JSPS KAKENHI grant (JP24K22305, https://kaken.nii.ac.jp/en/grant/KAKENHI-PROJECT-24K22305/) from the Japan Society for the Promotion of Science, JST CREST (JPMJCR22D2) and JSPS KAKENHI grant (JP25K03084, https://kaken.nii.ac.jp/en/grant/KAKENHI-PROJECT-25K03084/). Illustration of the mouse brain silhouette used in this manuscript was sourced from Sci-Draw, with thanks to Roberta Schellino (https://scidraw.io/drawing/646). We acknowledge Sci-Draw for making this valuable resource openly available. We would like to thank Prof. Toshio Aoyagi for fruitful discussion and comments about estimating the stationary distribution of phase difference in the phase-coupled oscillators using Fokker-Plank Equation.

## Supplementary Material

### Supplementary Results

In the main manuscript, we proposed a method to extract causal directed network structures from observed neural oscillatory signals by combining data-driven dynamical network modeling (Yokoyama & Kitajo, 2022) with a time-series causal discovery (CD) method (Hyvärinen et al., 2010; Shimizu, 2014). Our proposed method consists of three steps: (1) data-driven modeling of dynamical network systems (i.e., estimating ODEs), (2) extracting a directed information pathway that explains the causal DAG from ODE systems using our proposed method, and (3) evaluating causal effects and their causal ordering to determine causal hierarchical structures in the systems. To bridge the first step to the second, we proposed an approximation approach that maps the ODE estimated in step (1) into a structural vector autoregressive (SVAR) model with directed acyclic causal structures, using the Jacobian matrix and Padé approximation (Bergstrom, 1984; Oud et al., 2018; Oud & Delsing, 2010; Voelkle & Oud, 2013). In our method, the estimation algorithm for this linearization is established by incorporating Jacobian-matrix information into a vector autoregressive linear non-Gaussian model (VAR-LiNGAM), which is used as the estimation algorithm in SVAR-based time-series causal models. By doing so, we can transform ODE-based dynamical systems into SVAR-based time-series causal models in a data-driven manner (this method is termed j-VAR-LiNGAM; Jacobian-informed VAR-LiNGAM). The method is validated by showing that it enables causal hierarchical ordering in the observed synchronous neural dynamics reflected in both numerical and empirical data (see Figs. 3--8 in the Results section of the main manuscript).

This text provides the supplemental results of Simulations 1 and 2 in the main manuscript. As mentioned in the main manuscript, to estimate the Jacobian-informed linearized SVAR model using j-VARLiNGAM, our method requires determining the Jacobian matrix in estimated ODEs around the steady-state solutions obtained by solving Fokker-Planck equations (FPE) (Ota et al., 2020). This text shows the results of determining the steady-state solutions of estimated ODEs for each numerical simulation.

#### S1. Additional result in Simulation 1

The results of solving FPE in Simulation 1 are shown in Fig. S1. In this simulation, we confirmed how the estimation accuracy of causal structures in our method is affected by the strength of the synchronous dynamics in the dynamical network systems. To test this, we applied our method to synthetic phase observations generated by the Kuramoto-like phase-coupled oscillators with various conditions of global coupling parameters. This approach allowed us to extract causal hierarchical structures from the observed phase data, particularly when the systems were in weakly coupled limit cycle states prior to phase transitions.

In this simulation, the global coupling strength is a control parameter of phase transition in network systems for generating synthetic observations. To focus on the effect of this parameter for estimating steady-state solution of estimated ODEs that is required to obtain the Jacobian matrix for j-VARLiNGAM, we confirmed the results in solving FPE in the ODEs and compared the results depending on this parameter setting of Fig. S1A and B shows the results in *K* = 0.4 (i.e., weakly coupled limit cycle states) and *K* = 4.0 (i.e., almost perfect synchronous state), respectively. In this simulation, we can obtain the steady-state distribution of phase difference among all pairs of oscillators, by solving FPE (see Eqs. (32-35)). As can be seen in Fig. S1 and B, the estimated distribution of (i.e., solution of FPE) is well fitted to the empirical distribution for each pair of oscillators for both parameter settings: *K* = 0.4 and *K* = 4.0. This result supports that our estimated Jacobian matrix of ODE captures the valid phase interaction in the observed data. Combining the results in Fig.S1 and Fig. 3 in the main manuscript, our method can correctly extract directed acyclic causal structures based on the dynamical property reflected in the Jacobian matrix determined from an accurate solution of FPE in ODEs. To guarantee the identifiability and accuracy of our proposed method, both accurate estimation of the Jacobian matrix and the causal model should be satisfied. The results suggest that both requirements are satisfied.

**Figure S1:**
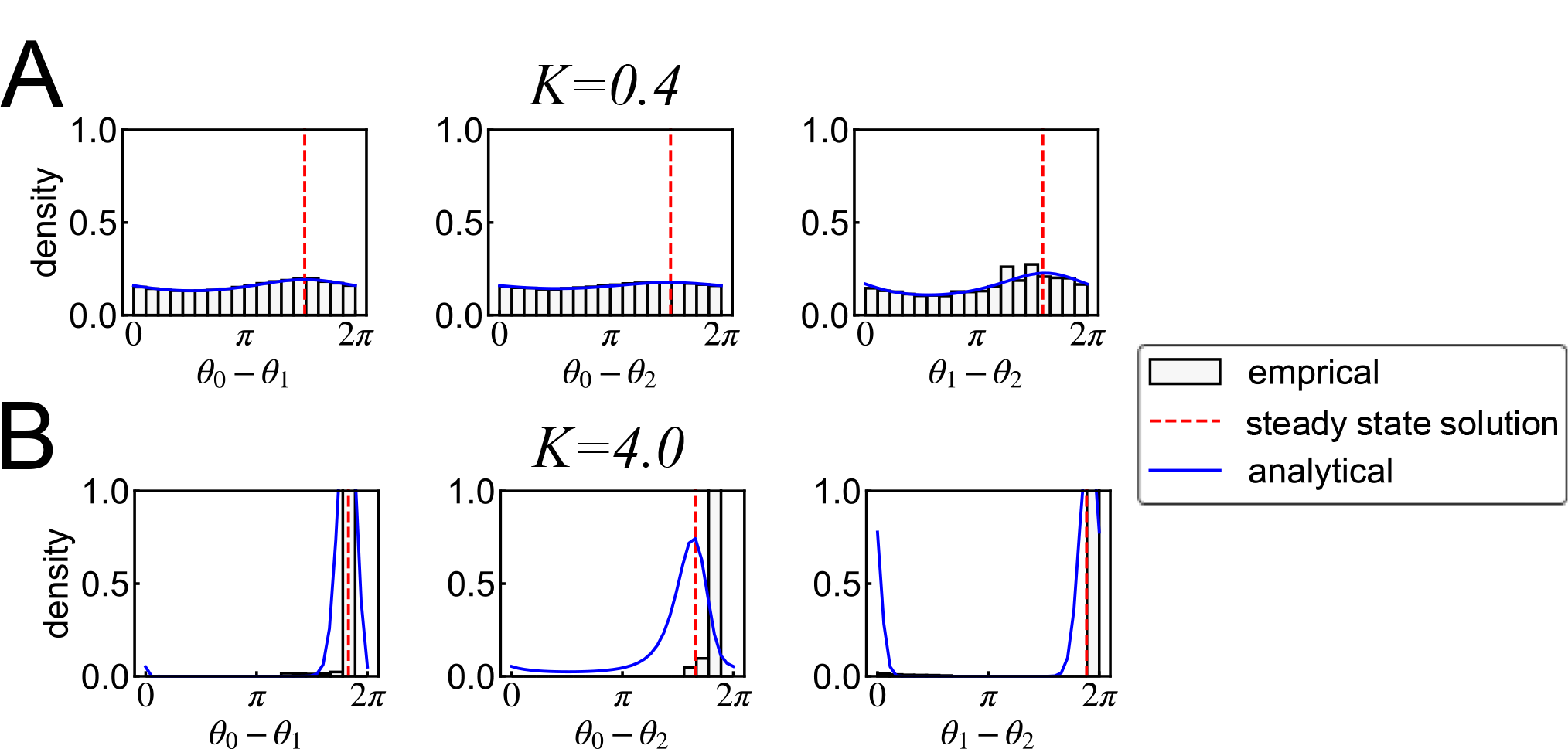
Comparisons of steady state distribution of phase differences in the estimated phase-coupled oscillators using Fokker-Plank equation (FPE) (Simulation 1). (A, B) Estimated distributions of phase difference Ψ_*ij*_ for each pair of oscillators, *i, j*, with conditions under *K* = 0.4 and *K* = 4.0. Gray histogram indicates the empirical distribution of Ψ_*ij*_. Blue bold and red dotted lines indicate the estimated distributions and steady state solution of Ψ_*ij*_ using FPE.

#### S1. Additional result in Simulation 2

This section provides the additional results in Simulation 2 of the main manuscript. In this simulation, we applied the synthetic phase data generated from a phase-coupled oscillators model that contains more complex phase interacting dynamics among oscillators. As can be seen in Fig. 6 of the main manuscript, our method can correctly detect both causal structures and hierarchical ordering in oscillations, even when the other conventional method (standard VAR-LiNGAM; Hyvärinen et al (2010) and PCMCI; Runge et al (2023)) struggles for determination (see main text for more details).

This section shows the results of estimating the steady-state solution, which is required to determine the SVAR-based linearized time-series causal model using j-VARLiNGAM (Fig. S2). As shown in Fig. S2, the estimated distribution of Ψ_*ij*_ is well fitted to the empirical distribution for each pair of oscillators, as well as Fig. S1. Moreover, each steady-state solution of were found as the peak of the estimated distribution Ψ_*ij*_. This result supports that our estimated Jacobian matrix of ODE captures the valid phase interaction in the observed data, and this fact indicates that the causal analysis results obtained from our proposed method, j-VAR-LiNGAM, in this simulation, shown in Fig. 6 of the main manuscript is reliable.

**Figure S2:**
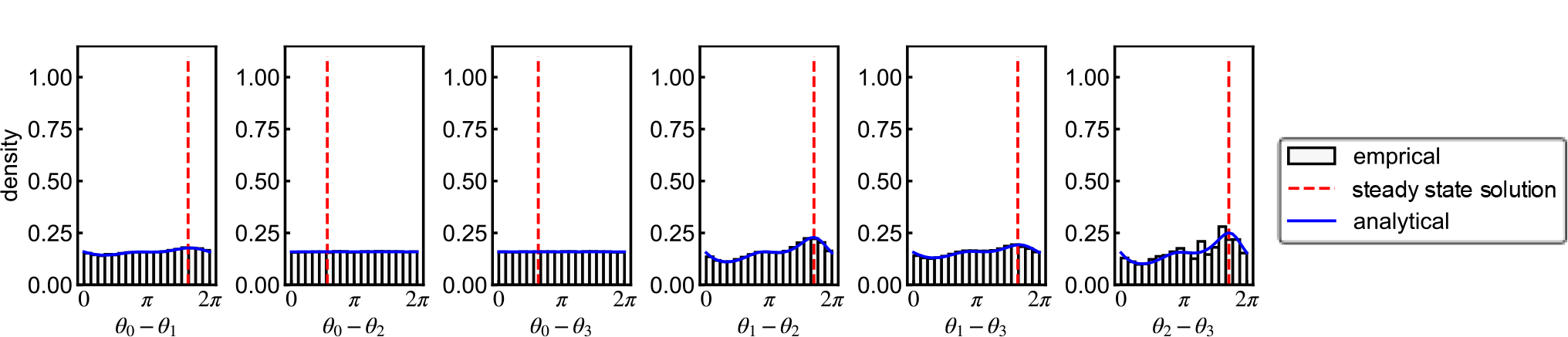
Result of steady state distribution of phase differences in the estimated phase-coupled oscillators using Fokker-Plank equation (FPE) (Simulation 2). It shows the estimated distributions of phase difference Ψ_*ij*_ for each pair of oscillators, *i, j*. Gray histogram indicates the empirical distribution of Ψ_*ij*_. Blue bold and red dotted lines indicate the estimated distributions and steady state solution of Ψ_*ij*_ using FPE.

